# Knowledge, attitudes and practices (KAPs) and risk factors of brucellosis at the human-animal interface in the Nile Delta, Egypt

**DOI:** 10.1101/607655

**Authors:** Ekram W. Abd El-Wahab, Yamen Hegazy, Wael F. El-Tras, Ashraf Mikeal, Ahmed F. Kapaby, Mahmoud Abdelfatah, Mieghan Bruce, Mahmoud M. Eltholth

## Abstract

**Background:** Brucellosis is a highly contagious zoonosis affecting human and almost all domestic species. It is a multi-burdens disease leading to severe economic losses due to disability in humans in addition to abortion, infertility and reduced milk production in animals. An Important element for effective prevention and control of brucellosis is to improve knowledge, attitude and practices of the community.

**Objective(s):** This study aimed to assess the knowledge, attitudes and practices (KAP) related to brucellosis at human-animal interface and to determine the risk factors for human infection in the Nile Delta, Egypt.

**Methods:** A matched case-control study was conducted at the main fever hospitals located in 6 governorates in the Nile Delta, Egypt between June 2014 and June 2016. Face-to-face interviews with cases and controls was done using a structured questionnaire. Differences in proportions of KAP variables among the cases and controls were evaluated by Pearson’s Chi square test and a *p* value <0.05 was set as a level of significance. A multivariate conditional logistic regression analysis was built to determine the risk factors for Brucella spp. infection among study participants.

**Results:** A total of 217 cases and 434 controls matched for age, gender and sociodemographic characteristics were enrolled and interviewed. In total, 40.7% of the participants owned animals in their households and lived in shared accommodation with animals [48.8% of cases vs 36.9% of controls; (*p*= 0.003)]. The majority (78.1%) used to accommodate cows and buffaloes with sheep and goats. Human brucellosis cases experienced more animal abortions comparing to the controls [(23.5% vs 9.7%, respectively), (*p*= 0.0003)]. The majority of the participants (82.4%) did not notify authorities in case that abortion occurs in their owned animals. Apparently, 67.4% of the participants [(70.0% of the cases vs 66.1% of the controls) (*p* = 0.315)] had not ever heard about brucellosis. The overall mean practice score regarding animal husbandry, processing and consumption of milk and dairy products was significantly lower among cases comparing to controls (−12.7±18.1 vs 0.68±14.2 respectively; *p*<0.0001). Perceived barrier for notification of animal infection and/or abortion was significantly higher among cases (*p*= 0.034) and positively correlated with participants’ education. Results of univariate analysis showed that participants who have animals’ especially small ruminates were at a higher risk of getting *Brucella* spp. infection than others. In the proposed multivariate conditional logistic regression model, the predictors of having brucellosis infection were consumption of unpasteurized milk, having consumed dairy products in the last 3 months before the study, consumption of yoghurt or home-made cheeses and involvement in contact with animals [OR (95% CI) = 4.12 (1.62 - 10.75); 2.71 (1.06 – 6.93); 2.51 (1.21 – 5.24); 1.96 (1.17- 3.30), *p*<0.05; and 4.97 (2.84 - 8.72)], respectively. Participants who take more protective measures against infection were at a significant lower risk of being diseased with brucellosis; [OR (95% CI) = 0.23 (0.10 - 0.58); *p*<0.001], respectively. A model predicting risk factors for brucellosis among those who own animal showed that frequent abortions per animal increased the chance for brucellosis infection among human cases by 49.33 fold [(95% CI)= (8.79 – 276.91); *p*= 0.001] whereas the practice protective measures with animals was protective for humans as well [OR (95% CI)= 0.11 (0.03-0.45); *p*= 0.002].

**Conclusion:** Consumption of dairy products stands side by side with the contact with infected animals particularly aborted ones as the major risk factors for *Brucella* spp. infection among humans in Egypt. On the other hand, there is a poor knowledge, negative attitudes and risky behaviors among villagers which increase the magnitude of the risk of brucellosis transmission at the human-animal interface. This supports the need for integrating health education in the national brucellosis control programs in Egypt with a special emphasis on hygienic animal husbandry, disease notification and benefits of animal vaccination.

**Author summary:** Zoonotic brucellosis has a vast global burden and remains neglected in many areas of the world despite notable advances in disease containment strategies. Despite the implementation of a national brucellosis control program in Egypt, the challenges for the disease eradication are intractable and multifaceted. We modeled in the present study the multivariate factors for brucellosis persistence in Egypt which apparently pointed to lack of basic understanding of the nature of brucellosis, traditional practices, beliefs and risky behaviors being undertaken on farms and at households across a wide region of the country. Predominantly, consumption of dairy products from unregulated sources; underreporting animal infection and abortion; underutilization of animal vaccination service; unsanitary disposal of abortus; use of milk of infected/aborted ruminants and lack of protective measure when practicing animal husbandry. Together, these conflict with disease intervention strategies and contribute to disease spread and re-emergence. The proposed model can provide a framework for future containment strategies that should be adopted to support and enhance the adherence to the current national brucellosis control program.

## Introduction

Brucellosis is a neglected zoonosis of public health and economic significance in most developing countries. Although the disease is well controlled in many countries, it remains endemic in others with highest records in the Middle East and central Asia [1,2] In most of these countries, the primary source of human infection is cattle, buffaloes, sheep and goats infected with *Brucella* spp. [3,4]. Therefore, measures and strategies aimed to reduce the prevalence of brucellosis in animals are considered the most effective means of controlling human infection [5].

In animals, the disease is highly contagious affecting almost all domestic species, leading to severe economic losses due to abortion, infertility, loss of milk production and cost of veterinary care. Although, reliable estimates of the frequency of brucellosis among ruminants in Egypt are lacking due to inability to test all eligible animals periodically and properly [6]. Recent studies for the prevalence of brucellosis in ruminants indicate that the disease is endemic in all ruminant species with high prevalence mounting to 26.6% [7–12]. These findings question the efficacy of the applied national control programme for brucellosis which was established in the early 1980s and implies on serological surveys (test and slaughter, and compensation are paid for livestock owner), milk ring testing for pooled milk, and voluntary vaccination of ruminants [11,13].

Human infection with brucellosis has been reported in different studies in different geographical areas. The median number of foodborne illnesses, deaths, and Disability Adjusted Life Years (DALYs) were reported by the WHO in 2010 to be 832,633 (95% CI=337,929–19,560,440); 4,145 (95% CI=1,557–95,894); 264,073 (95% CI=100,540–6,187,148) [2]. In Egypt, the rate of human infection is greatly affected by the rate of disease in animals [13–16]. Direct contact with infected animals, aborted foeti, foetal membranes, and vaginal discharges of infected animals are risk factors for human infection with brucellosis. Further, humans can be exposed to infection through ingestion of un-pasteurised milk and raw milk products, such as soft cheeses and yogurt, which are commonly consumed in Egypt. Establishing the relative contribution of occupational and food-borne risk factors will inspire more targeted public health programmes. In this context, Jennings et al (2007) recommended that further studies are needed to assess the risk of human exposure to brucellosis via different exposure routes [15].

So far in Egypt, there is no specific studies which had tackled the economic and logistic causes for the failure of the control programme on small livestock holders and on the national level as well. However, Holt et al., reported that one of the major causes of the failure of the national brucellosis control programme in Egypt is the lack of compliance of the farmers with this programme due to the insufficient compensation for the slaughtered infected/aborted ruminant which is usually the key incentive used for test-and-slaughter [10]. Therefore, the aims of this study are to explore the risk factors of *Brucella* spp. infection among humans and to study the specific KAP components that contribute to the poor response to brucellosis control at the human-animal interface in the Nile Delta (Egypt). Further, we seek to point out the critical points which need to be managed and addressed in future research. This will build baseline data to design a framework for identifying problems facing the current national brucellosis control programme in smallholders and the nation wise; besides helping scientists and policy makers in developing the appropriate control strategy.

## Methods

### Study design, setting and population

A case-control study was conducted between June 2014 and June 2016 in the main fever hospitals in the Nile Delta region. Cases and controls were identified from the following governorates; Alexandria, El-Behira, Gharbia, Kafrelsheikh, Dekahlia and Matrouh (Figure 1). These governorates have a high density of people and livestock, where human and animals are living in close proximity, particularly in small-scale farming system. Cases were included in the study once they are identified and controls were sampled over the same time period. Previous studies in the area identified that risk factors [adjusted odds ratio (AOR)] of having brucellosis are: having sheep (6.2), being a farmer or butcher (4.5), having aborted animal (3.5) and increasing age 1.04 per year [17].

**Figure 1:**
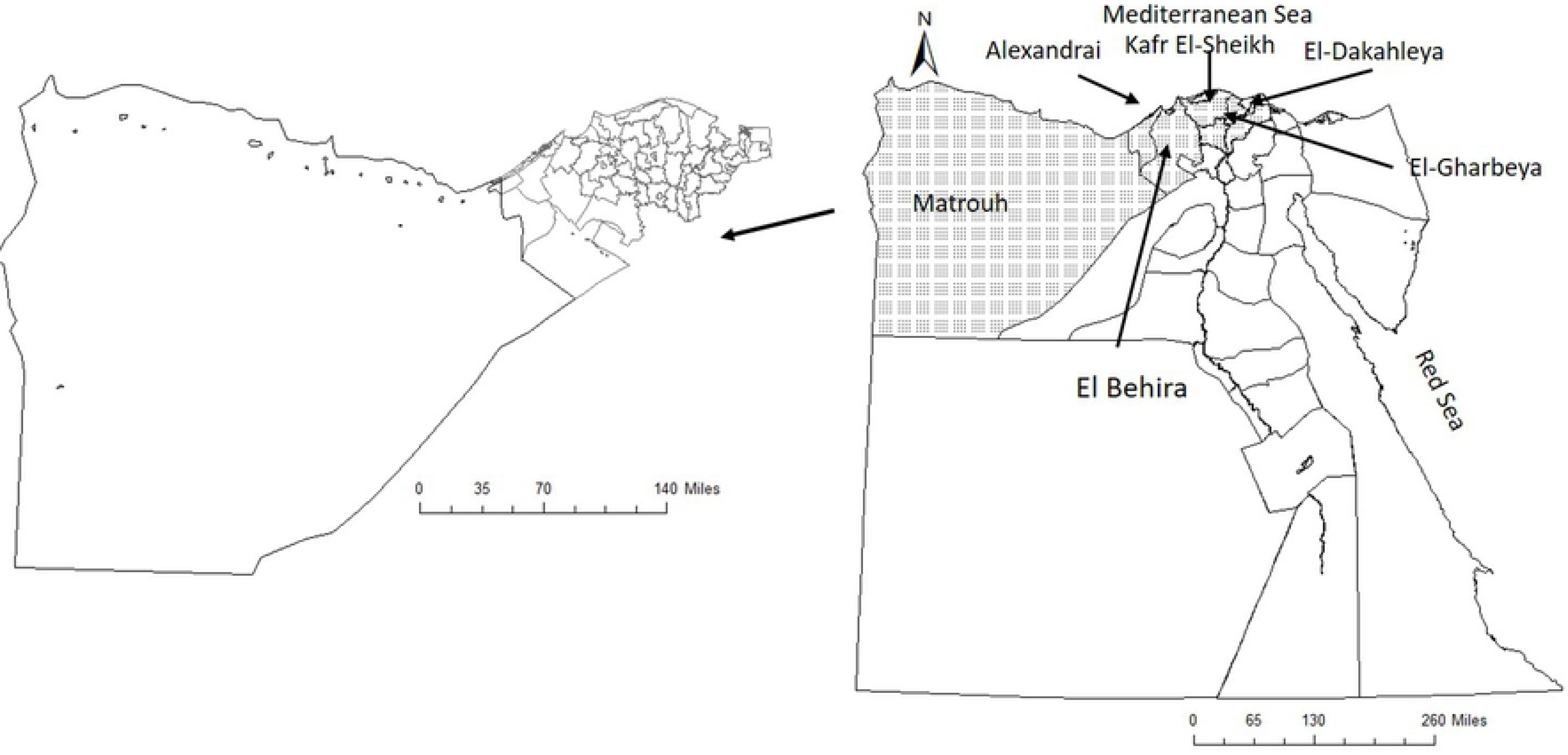
A choropleth map of Egypt showing the administrative boundaries of the governorates: the dotted governorates represent the study area. The map was created using Quantum GIS (Quantum GIS Development Team 2017)

The sample size was calculated using Win Episcope 2.00 for a matched case-control study based on 80% power and 95% confidence interval with 40% estimated exposure rate for controls and 2.2 minimal OR to be significantly detected. The minimal required sample size was 206 (103 cases and 103 controls). However, we included in the study 217 cases and the number of controls was doubled to have 434 controls. Cases were defined as individuals seeking medical care at the fever hospitals within the study area. The case definition of the World Health Organisation (1999) was applied: “Cases clinically presented with high fever, profuse night sweating, fatigue, anorexia, weight loss, headache and arthralgia”. The diagnostic work-up of suspected cases included Rose Bengal Test (RBT) and the positive case were confirmed by Standard Tube Agglutination Test (SAT) with titre > 1:160). To reduce selection bias, for each enrolled case, two matched controls for age, gender and residency (rural or urban) were selected from persons seeking medical care for other health conditions at the same hospitals. Controls were confirmed as serologically free of brucellosis using SAT with titre > 1:160.

No incentives we provided to participate in the study

### Data collection

a structured questionnaire was used for collecting sociodemographic (for the study population and other family members in the same houses of the study population) and epidemiological data on potential risk factors for an individual being seropositive against brucellosis. This included dairy product consumption habits, animal husbandry practices and history of exposure of humans and animals to brucellosis in the same household. Further, information on the collaboration with health service in case of human or animal infection with brucellosis was gathered. The knowledge, attitudes and practices relating to brucellosis were assessed through the use of open questions (S1 File). The questionnaire was developed, pre-tested and validated to have a good insight in the small-scale dairy farming sector in the study area. All interviewers were trained to standardise the interviewing method. During the interviews, the questions were continuously evaluated to make sure that the farmers understood them correctly.

### Calculation of scores

Knowledge was assessed through 5 open questions. Correct complete answers were scored 3 points, incomplete answers were scored 1 point and wrong/do not know answers were scored 0 point. Attitude (perceived barrier, perceived risk, perceived susceptibility) were assessed through 7 open questions. For perceived barriers, 1 point was given for each response. For perceived risk and perceived susceptibility, responders were asked to rate their responses as none, low, mild, moderate and high. Accordingly, responses were rated on a 5-point Likert scale [0 low-5 high]. Hygienic practices were assessed via 26 practice statements measured by yes/no answers or through a three-point frequency rating with the options “always”, “sometimes” and “never”. Safe practices were given score of 1 and risky ones were scored −1. The total score was graded as “good practice” if exceeded 75% of the total score, and poor practice for scores< 50% to −100%. Milk consumption score was calculated as number of days of milk consumption per a person in a week; 0= no consumption, 1= consumed 1-3 times in a week, 2= consumed >3 times in a week.

### Statistical analysis

Data were analysed using SPSS software version 18.0 (SPSS Inc., Chicago, IL, USA) and SAS 9.2 (SAS Institute Inc 2008). Differences in proportions among cases and controls were evaluated by Pearson’s Chi-square test. Differences in the total mean KAP scores were analysed using student t-test. A *p* value < 0.05 was set as a level of significance. The association between the potential risk factors and a brucellosis case was examined using a multivariate conditional logistic regression model, with individual status as a control or a case as the response variable. The selection of variables to be included in the multivariate model was carried out in two steps. Initially, a univariate logistic regression model was built to determine the association between each of the examined variables and disease status of each individual where variables for which *p*> 0.2 were excluded from further analysis. The collinearity between pairs of variables with a *p*< 0.05 in the previous step was assessed by calculating the Phi correlation coefficient. The significance of this collinear association was examined using chi square test. In the case of a pair of variables with a significant association (*p* < 0.05), the variable judged as the most biologically plausible was used as a candidate in the multivariate analysis. All variables passed the previous 2 steps were incorporated in the final multivariate conditional logistic regression model. A manual stepwise selection approach was used for the selection of variables in that model to keep only variables with *p* < 0.05 in the final model. All two-way interactions between variables retained in the model were assessed. Testing for confounder was carried out by monitoring the change of logit of factors by removing a suspected factor from the model.

### Map creation

An electronic map of Egypt was provided by the General Organization of Veterinary Services (GOVS) in Egypt. A choropleth map was built for the geographic distribution of different study locations within Egypt using Quantum GIS (Quantum GIS Development Team 2017) www.qgis.org.

## Results

### 1. Socio-demographic characteristics of the study population

The study comprised 217 cases of human brucellosis and 434 matched controls with an overall mean±SD age of 35.2±13.9 and 35.6±14.5 years, respectively. There was no statistically significant difference between cases and controls regarding their socio-demographic characteristics including the occupation, education and income (Table 1). The number of reported household members of *Brucella* spp. infected cases in the 3 months before to the study was significantly higher than those households’ members of controls [(4.6% *vs* 1.8%) (*p*=0.015)] (Table S1). Details about the sociodemographics of the household members of the study population are portrayed in (Table S1). The results showed also that infected cases and their household members sought medical advice in private clinics (96.1%) and 96.8% of them were then referred to fever hospital for admission. About 95% of infected participants used to get medicine from the pharmacy as a first step before visiting the doctor.

**Table 1:**
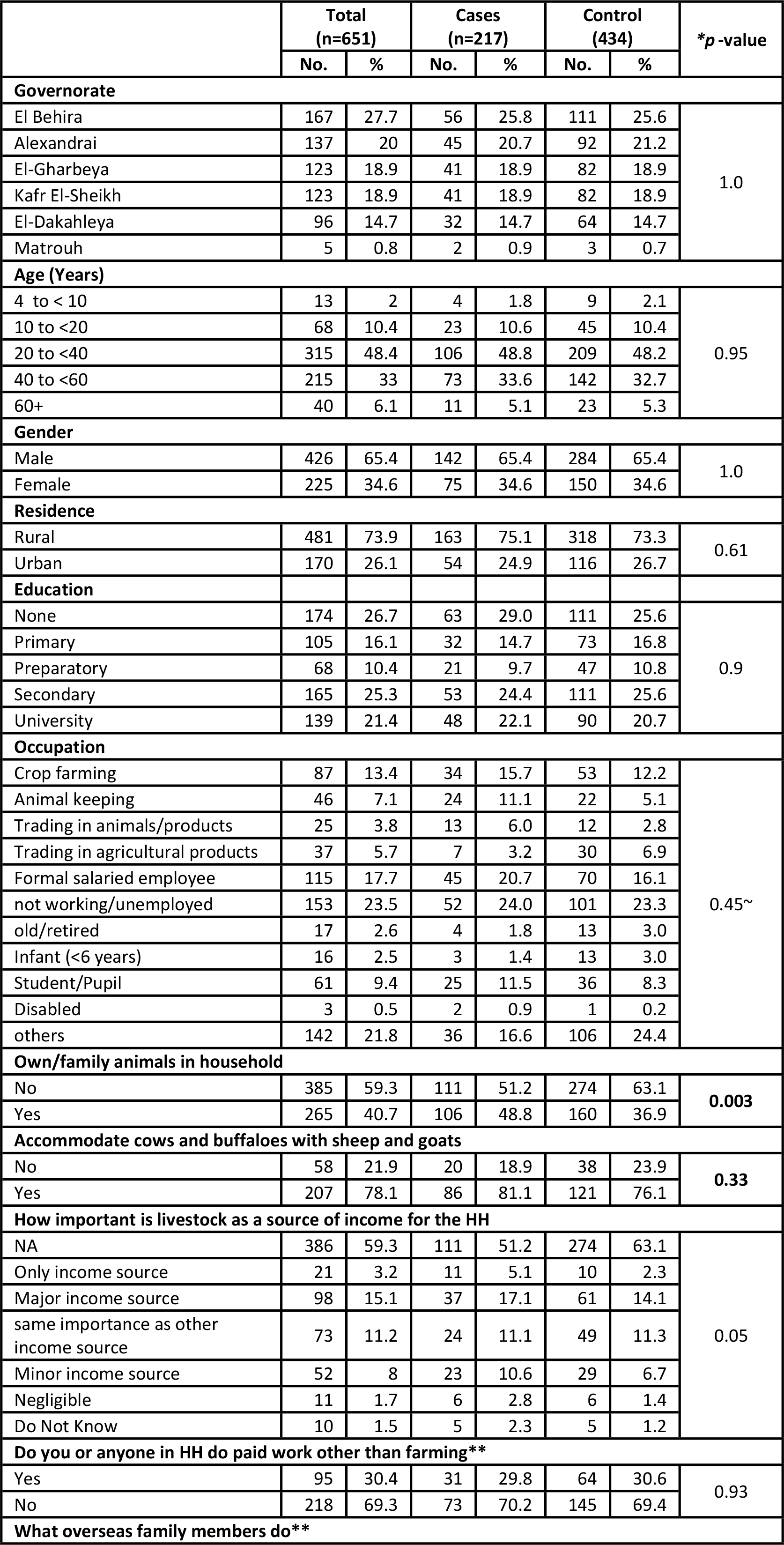
Sociodemographic characteristics of the study population

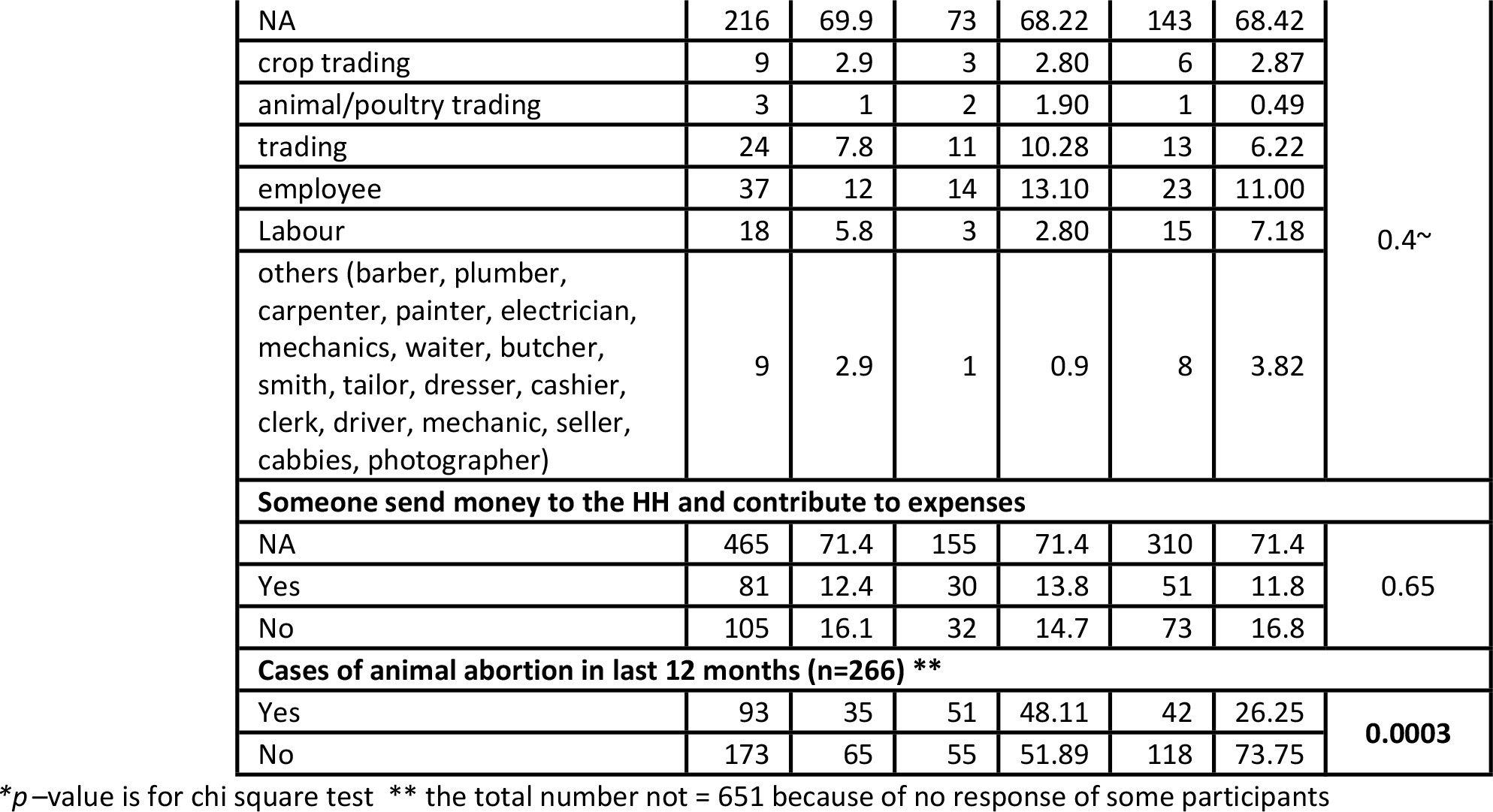

### 2. Owned animals in household and exposed to aborted animals

In total, 40.7% of the participants owned animals in their households [48.8% of cases *vs* 36.9% of controls; (*p*=0.003)]. The owned animal species included cows (91.7%), buffaloes (74.1%), sheep (62.8%), goat (41.0%), and donkeys, camels or both (35.7%), the proportions of different animal species belonged to both cases and control participants are shown in Figure 2. The number of sheep and goats owned by households was significantly higher among cases comparing to controls (*p* < 0.0001). The majority of livestock owners (78.1%) used to accommodate large (cows and buffaloes) and small ruminants (sheep and goats) together with no significant difference between cases and controls (*p* = 0.33) (Table 1).

**Figure 2:**
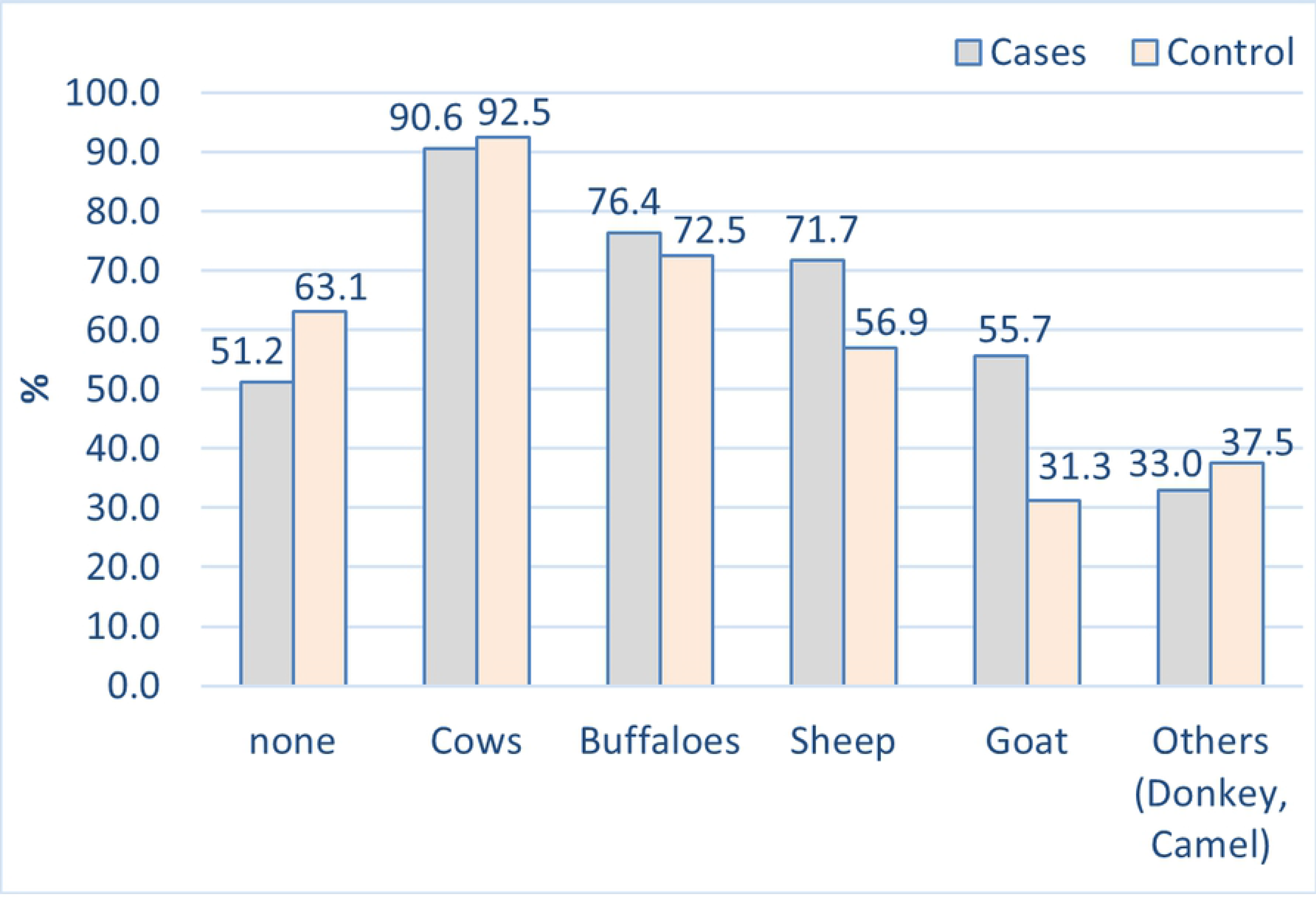
Percent of owned animal species in households of cases and controls

Human brucellosis cases were more likely to experience occurrence of animal abortions in the 12 months preceding the occurrence of human infections comparing to the controls [(23.5% *vs* 9.7%, respectively), (*p*=0.0003)] (Table 1). All of different ruminant species owned by participants experienced abortion incidents in the 12 months preceding the occurrence of human infections (Table S2). The most number of participants claimed that they do not know the cause of abortion with (Table S2). Livestock owners either cases or controls tend to call private veterinarian (83.2%) more than governmental veterinarian (12.8%) when having abortion (Table S3). Biological samples were collected to understanding the reasons for abortion from 10.9 % only of the incidents (Table S3) with no significant difference between controls and cases in such practice (*p*= 0.34). These samples were blood samples (82.4%) and foetal membranes/tissue/organs from aborted foeti (11.7%) or vaginal swab (5.9%). Outcomes of the biological sample analysis for different animal species as being positive or negative for brucellosis are summarized in (Table S3), the proportion of seropositive aborted cows and goats to brucellosis belonged to human cases was significantly higher than those among control participants (Table S3). The majority of the participants (82.4%) did not notify authorities in cases of abortion in the owned household animals and only 10.6% did with no significant difference between cases and control participants in that (Table S3). Reasons for not notifying animal abortion are listed in (Table S3). Most of the participants admitted that they do not know that they have to notify that incidents while some of them worry that health authority slaughters the diseases animals and not give sufficient amends (Table S3). Data about how participants handle aborted animals, dispose aborted foeti and process milk from aborted animals is displayed in (Tables S4 and S5). Most of them keep aborted animals for fattening or reproduction or sell them for reproduction and throw the foetal membranes and aborted foeti in water canals. About half of participants consume or sell for consumption their milk and dairy products either with or without heat treatment.

A total of 6 (2.3%) human cases and none of the controls (0.0%) declared that they had brucellosis infected animal 12 months before the study as proved by private laboratory investigations (50.0%), or across the national brucellosis control campaigns (33.3%) (Table S6). Causes of rejecting the notifications and what did national authorities do with the confirmed animal cases are summarized in table S6. One of the main causes of rejecting notification is that the participant does not know that he has to notify. If the infected ruminant was lactating, half of participant declared that they throw the milk away. On the other hand, for those who use the milk of infected animals, usage of heat treated milk, selling raw milk and processing homemade cheese, cream, butter, and ghee are practised with no significant difference by both cases and controls (Table S7).

The number of control participants followed protective measures for protecting their owned household ruminants from infection with *Brucella* is significantly higher than number of cases participants (Table 2). Among these protective measures, vaccination (36.5%) was the most commonly reported, followed by regular checkup at veterinary clinic (27.1%), cleaning animal house and keeping it well ventilated (15.8%), and treatment of infected ruminants (10.5%). Other measures are listed in (Table 2). The practice and the positive attitude towards animal vaccination were significantly higher among controls comparing to cases (*p*<0.0001) (Table S8).

**Table 2:**
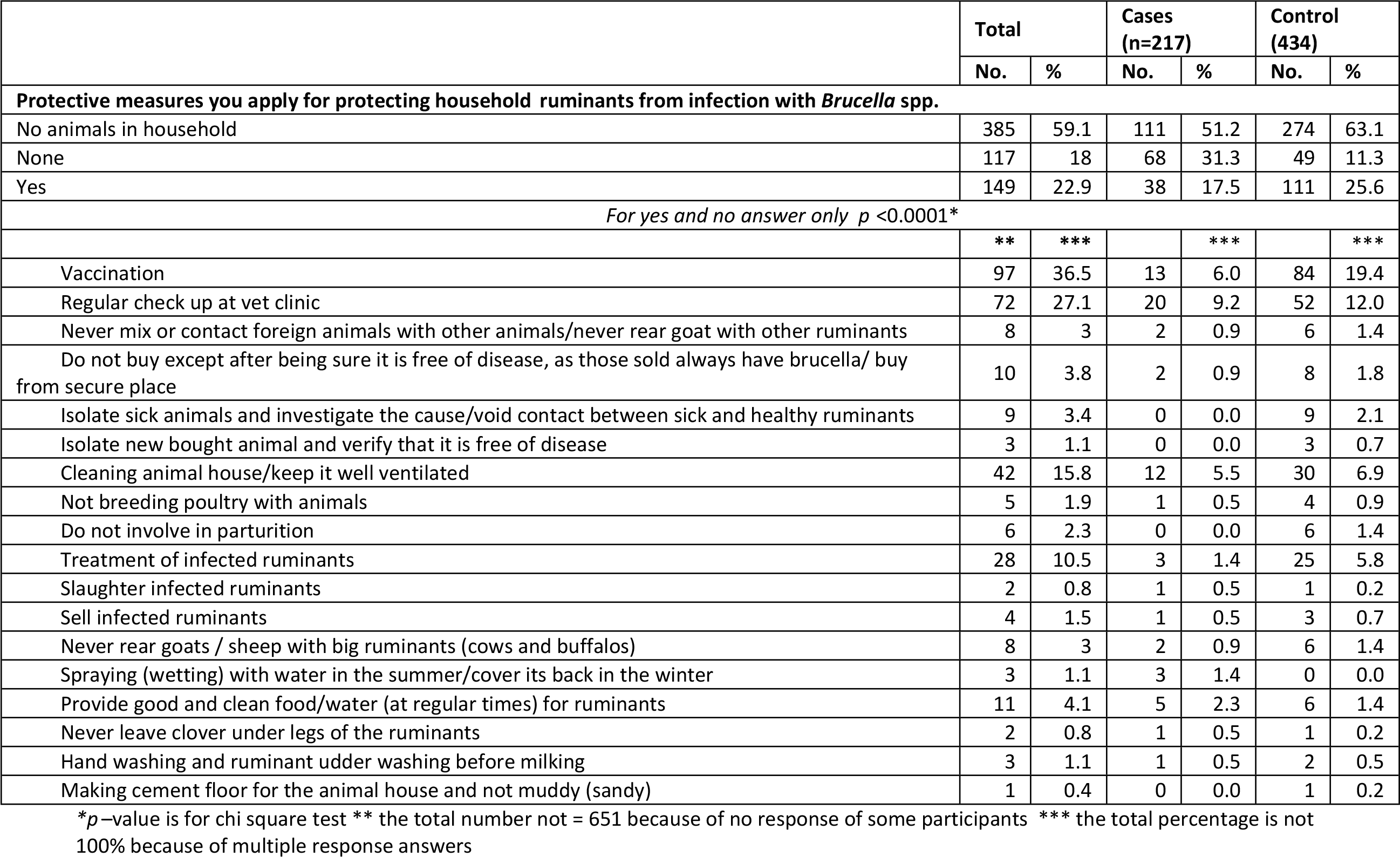
Protective measures applied by cases and controls for protecting household ruminants from infection with Brucella spp.

Interestingly, cases admitted the adoption of some measures for protecting household members from infection with *Brucella* spp. more than controls [54.8% of cases vs 29.3% of controls; *p*<0.0001]. These included boiling milk [30.9% of cases vs 28.3% of controls], buying pasteurised milk [14.3% of cases vs 2.1% of controls], not involving in parturition/parturition of infected animals [14.3% of cases vs 5.5% of controls], and using personal protective equipment against occupational hazards [6.0% of cases vs 4.6% of controls]. On the other hand, vaccinating animals was more frequently stated by controls [1.8% of cases vs 8.3% of controls] (Table 3).

**Table 3:** Protective measures applied by cases and controls for protecting household members from infection with Brucella spp.

|  | Total |  | Cases (n=217) |  | Control (434) |  |
| --- | --- | --- | --- | --- | --- | --- |
|  | No. | % | No. | % | No. | % |
| **Protective measures you apply to protect yourself and household members from brucella infection** |  |  |  |  |  |  |
| None | 405 | 62.2 | 98 | 45.2 | 307 | 70.7 |
| Yes | 246 | 37.8 | 119 | 54.8 | 127 | 29.3 |
| $p < 0.0001^*$ | | | | | | |
|  | ** | *** |  | *** |  | *** |
| Animal vaccination | 40 | 6.1 | 4 | 1.8 | 36 | 8.3 |
| Boiling milk | 190 | 29.2 | 67 | 30.9 | 123 | 28.3 |
| Do not involve in parturition/parturition of infected animals | 55 | 8.4 | 31 | 14.3 | 24 | 5.5 |
| Do not involve in disposing placenta | 2 | 0.3 | 0 | 0.0 | 2 | 0.5 |
| Regular animal checkup | 14 | 2.2 | 1 | 0.5 | 13 | 3.0 |
| Let vet doctor responsible for parturition and abortion | 7 | 1.1 | 0 | 0.0 | 7 | 1.6 |
| Animal slaughter if infected | 2 | 0.3 | 1 | 0.5 | 1 | 0.2 |
| Do not involve in abortion | 8 | 1.2 | 0 | 0.0 | 8 | 1.8 |
| Do not be very close to ruminants (breathing) | 3 | 0.5 | 2 | 0.9 | 1 | 0.2 |
| Do not drink milk/eat homemade cheese or fermented milk of infected ruminant/buy from secure source | 24 | 3.7 | 9 | 4.1 | 15 | 3.5 |
| Treatment of infected ruminants | 10 | 1.5 | 0 | 0.0 | 10 | 2.3 |
| Not coming in contact with infected ruminant | 19 | 2.9 | 6 | 2.8 | 13 | 3.0 |
| Sell infected ruminant | 2 | 0.3 | 0 | 0.0 | 2 | 0.5 |
| Using personal protective equipment against occupational hazards/wearing gloves, long boot | 33 | 5.1 | 13 | 6.0 | 20 | 4.6 |
| Not drinking goat milk | 2 | 0.3 | 1 | 0.5 | 1 | 0.2 |
| Personal cleaning/cleaning animal house, applying sand on ground of animal house | 29 | 4.5 | 6 | 2.8 | 23 | 5.3 |
| Separate animal house from own house/avoid going to animal house | 3 | 0.5 | 1 | 0.5 | 2 | 0.5 |
| Do not rear goats and sheep with other ruminants (cows and buffalos) | 1 | 0.2 | 0 | 0.0 | 1 | 0.2 |
| Prepare/eat cheese and milk products from healthy ruminants | 4 | 0.6 | 0 | 0.0 | 4 | 0.9 |
| Avoid animal contact when having hand wound | 1 | 0.2 | 1 | 0.5 | 0 | 0.0 |
| Buying animals from secure source | 2 | 0.3 | 1 | 0.5 | 1 | 0.2 |
| Eating only milk and milk products from one's own ruminants | 1 | 0.2 | 0 | 0.0 | 1 | 0.2 |
| Being careful during handling poultry/ deal with animals | 5 | 0.8 | 1 | 0.5 | 4 | 0.9 |
| Cost of buying milk from secure source (neighbors you know their animal status) | 1 | 0.2 | 0 | 0.0 | 1 | 0.2 |
| Fighting flies and mosquitos | 3 | 0.5 | 0 | 0.0 | 3 | 0.7 |
| Provide clean clover, animal food, water | 2 | 0.3 | 1 | 0.5 | 1 | 0.2 |
| Avoid that children play with sheep | 1 | 0.2 | 1 | 0.5 | 0 | 0.0 |
| Do not rear animal at all | 3 | 0.5 | 1 | 0.5 | 2 | 0.5 |
| Buying pasteurized milk | 40 | 6.1 | 31 | 14.3 | 9 | 2.1 |
| Hand washing | 3 | 0.5 | 2 | 0.9 | 1 | 0.2 |
| Not take milk from aborted ruminant | 1 | 0.2 | 1 | 0.5 | 0 | 0.0 |

\* $p$ -value is for chi square test \*\* the total number not = 651 because of no response of some participants \*\*\* the total percentage is not 100% because of multiple response answers

Cases tended to involve more frequently in activities in which they come in contact with animals particularly when helping in animal parturition, abortion, disposing placental and aborted fetuses and slaughtering, (Table S9). Furthermore, cases were more frequently involved in dairy product processing (Table S10). Control participants were significantly higher than cases in selling/buying livestock products in markets and neighboring villages (Table S11). The proportion of cases consume small ruminates dairy products is higher than controls (Table S12), but the frequency of consumption of dairy products does not differ significantly between cases and control (Table S12).

Cases admitted the use of personal protective equipment such as gloves and mask and emphasized hand hygiene more frequently comparing to controls (Table S13).

### 3. Knowledge

More than two thirds (67.4%) of the participants had not heard about a disease called brucellosis with no significant difference between cases and controls [(70.0% of the cases *vs* 66.1% of the controls) (*p* = 0.315)] (Table S14). The source of knowledge was mainly through neighbours (74.6%), veterinarians (48.8%) and relatives (7.4%). About 82% of the participants did not know which animal species are more susceptible to brucellosis. Cows, sheep, goat, buffaloes, poultry/duck- and “all animal types” were mentioned by 17.4%, 17.1%, 13.5%, 11.4%, 3.1% and 2.8% of the respondents respectively (Table S14). Controls have a significant higher knowledge than cases participants that sheep, goats and cows are susceptible to *Brucella* spp. infection (Table S14). The age at which the animals are most susceptible to *Brucella* infection as stated by the study participants is displayed in (Table S14). Almost one third of the participants believe that *Brucella* can be transmitted to human with cases being more knowledgeable of that than controls (*p*<0.0001); 43.4% of cases *vs* 28.6% of controls (Table 4). The most frequently listed modes of transmission were drinking fermented milk/fermented milk of infected ruminant and contact/daily dealing with ruminants/infected ruminants (Table 4). Data about the perceived risk of the mentioned routes of transmission are displayed in (Table S15). In spite of cases being more knowledgeable than controls of the risk of consumption of infected dairy products and contact infected animals in transmission of *Brucella* spp. infection to human, there was significant lower knowledge among cases than control of the magnitude of the importance of these routs of infection as modes of brucellosis transmission to humans (Table S15).

**Table 4:** Knowledge among cases and controls regarding brucellosis and modes of transmission

|  | Total |  | Cases (n=217) |  | Control (434) |  |
| --- | --- | --- | --- | --- | --- | --- |
|  | No. | % | No. | % | No. | % |
| **Do you think that brucella can be transmitted to humans** |  |  |  |  |  |  |
| Yes, I think so | 219 | 33.6 | 95 | 43.8 | 124 | 28.6 |
| No, I do not think so | 235 | 36.1 | 60 | 27.6 | 175 | 40.3 |
| I do Not Know | 197 | 30.3 | 62 | 28.6 | 135 | 31.1 |
| **$p < 0.0001^*$** |  |  |  |  |  |  |
| **Mention route of transmission of brucella to humans (n=219)** |  |  |  |  |  |  |
|  | ** | *** |  | *** |  | *** |
| Do not know | 15 | 6.8 | 7 | 3.2 | 8 | 1.8 |
| Parturition | 25 | 11.4 | 6 | 2.8 | 19 | 4.4 |
| Animal abortion / of infected ruminant | 44 | 20.1 | 11 | 5.1 | 33 | 7.6 |
| Drinking milk of infected animal | 72 | 32.9 | 45 | 20.7 | 27 | 6.2 |
| Drinking unboiled milk | 78 | 35.6 | 19 | 8.8 | 59 | 13.6 |
| Help in getting placenta out | 4 | 1.8 | 1 | 0.5 | 3 | 0.7 |
| Drinking fermented milk/fermented milk of infected ruminant | 76 | 34.7 | 37 | 17.1 | 39 | 9.0 |
| Parturition of infected ruminants | 8 | 3.7 | 0 | 0.0 | 8 | 1.8 |
| Eating homemade cheese/cheese prepared from of infected ruminants | 83 | 37.9 | 42 | 19.4 | 41 | 9.4 |
| Contact/daily dealing with ruminants/infected ruminants | 87 | 39.7 | 52 | 24.0 | 35 | 8.1 |
| Breathing/animal odour | 14 | 6.4 | 4 | 1.8 | 10 | 2.3 |
| Accidental exposure to animal vaccine | 2 | 0.9 | 1 | 0.5 | 1 | 0.2 |
| Drinking unboiled goat milk | 1 | 0.5 | 1 | 0.5 | 0 | 0.0 |
| Exposure to animal/birds secretions/ cleaning poultry | 7 | 3.2 | 1 | 0.5 | 6 | 1.4 |
| Contacting goats/goat come near children food (smell it) and so infect them | 3 | 1.4 | 1 | 0.5 | 2 | 0.5 |
| By flies and mosquitos | 4 | 1.8 | 0 | 0.0 | 4 | 0.9 |
| By birds (poultry)/poultry odor/handling sick poultry/cleaning poultry | 5 | 2.3 | 1 | 0.5 | 4 | 0.9 |
| Through hand wound | 2 | 0.9 | 2 | 0.9 | 0 | 0.0 |
| Contaminated animal house/cleaning animal house | 5 | 2.3 | 1 | 0.5 | 4 | 0.9 |
| Exposure to ruminants' excreta (urine and stool)/ saliva | 5 | 2.3 | 2 | 0.9 | 3 | 0.7 |
| Having the animal house in the same own house | 2 | 0.9 | 0 | 0.0 | 2 | 0.5 |
| Handling raw milk and milk product | 1 | 0.5 | 0 | 0.0 | 1 | 0.2 |
| Milking infected ruminants | 1 | 0.5 | 0 | 0.0 | 1 | 0.2 |
| Passing into animal house without wearing long boots | 2 | 0.9 | 0 | 0.0 | 2 | 0.5 |
| Not washing hand after daily dealing with ruminants | 2 | 0.9 | 0 | 0.0 | 2 | 0.5 |
| Eating fatty meat | 1 | 0.5 | 0 | 0.0 | 1 | 0.2 |
| Eating dinner very late and direct sleeping after | 1 | 0.5 | 1 | 0.5 | 0 | 0.0 |
| If children play with sheep | 1 | 0.5 | 1 | 0.5 | 0 | 0.0 |
| Rearing cats and dogs | 1 | 0.5 | 0 | 0.0 | 1 | 0.2 |
| Rats | 1 | 0.5 | 0 | 0.0 | 1 | 0.2 |

\* $p$ -value is for chi square test \*\* the total number not = 219 because of no response of some participants \*\*\* the total percentage is not 100% because of multiple response answers

Cases were less knowledgeable about brucellosis disease and risk of infection and they reported more risky attitudes and practices (Table 5). The overall mean practice score was significantly lower among cases comparing to controls (−12.7±18.1 vs 0.68±14.2 respectively; *p* < 0.0001). Perceived barrier for notification of animal infection and/or abortion was significantly higher among cases (*p*=0.034) and positively correlated with participants’ education (Table 5). Knowledge was strongly associated with participants’ practice, perceived barriers, as well as perceived susceptibility and risk (Figure 3).

**Table 5:** Mean knowledge, attitude and practice scores among cases and controls, and by the level of education

| Score | Group |  | t- test | *p-value* | Education |  | t- test | *p-value* |
| --- | --- | --- | --- | --- | --- | --- | --- | --- |
|  | Cases | Control |  |  | Low Literacy | High Literacy |  |  |
|  | Mean±SD | Mean±SD |  |  | Mean±SD | Mean±SD |  |  |
| Knowledge | 4.3±7.2 | 4.8±10.6 | -0.8 | 0.418 | 4.9±9.7 | 4.4±9.5 | 0.55 | 0.58 |
| Practice | -12.7±18.1 | 0.68±14.2 | -9.5 | **0.0001** | -5.5±18.0 | -1.9±15.2 | -2.72 | **0.007** |
| Attitude | 3.4±7.6 | 4.6±11.0 | -1.7 | 0.093 | 4.5±10.0 | 4.0±9.8 | 0.59 | 0.56 |
| Perceived barrier | -0.3±0.7 | -0.2±0.64 | -2.13 | **0.034** | -0.28±0.75 | -0.12±0.53 | -3.25 | **0.001** |
| Perceived susceptibility | 1.5±3.2 | 1.9±4.0 | -1.2 | 0.24 | 1.6±3.24 | 1.9±4.1 | -0.82 | 0.414 |
| Perceived risk | 2.5±4.9 | 2.8±7.3 | -0.682 | 0.496 | 3.1±46.9 | 2.3±6.1 | 1.53 | 0.127 |
| Perceived susceptibility and risk | 4.0±7.0 | 4.7±10.5 | -0.961 | 0.337 | 4.7±9.6 | 4.2±9.3 | 0.73 | 0.47 |

**Figure 3:**
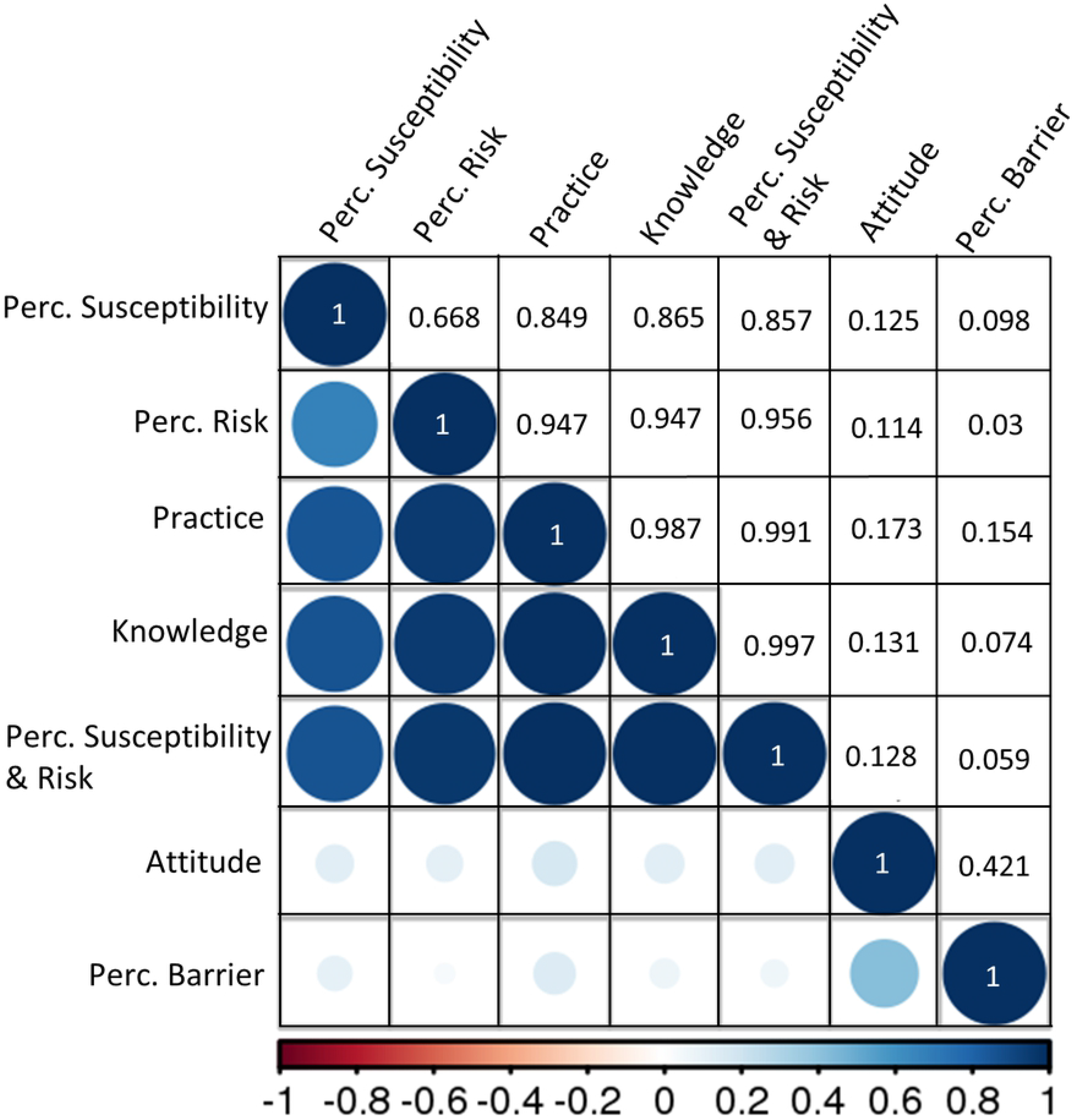
A corrplot visualizing a correlation matrix of the different KAP variables

### 4. Risk factor for human brucellosis infection

The results of the univariate relationships between independent variables and infection status of the participants are shown in (Table 6). Those who have animals in their houses especially small ruminants, non-vaccinated and aborted animals, participate in dairy products processing, consume different dairy products, do not follow protective measures to protect animals from infection, apply low number of protective measures to human from getting *Brucella* spp. infection and people who do not notify authorities for human or animal cases of brucellosis were at significant risk of getting brucellosis more than other participants.

**Table 6:** Results of a univariable model for the association between selected potential risk factors and individual Brucella spp. seropositive status

| Variable Categories |  | Number |  | OR | P < | 95% CI |
| --- | --- | --- | --- | --- | --- | --- |
|  |  | Case | Control |  |  |  |
| Have animals in the house | Yes | 106 | 160 | 2.7 | 0.001 | 1.68 – 4.5 |
|  | No | 111 | 274 | - | - | - |
| Number of animals in the house | 0 | 111 | 274 | - | - | - |
|  | 1-5 | 49 | 100 | 2.21 | 0.26 | 1.32- 3.70 |
|  | >5 | 57 | 60 | 8.2 | 0.001 | 3.88 – 17.32 |
| Have small ruminant in the house | Yes | 85 | 105 | 3.57 | 0.0001 | 2.1 – 6.03 |
|  | No | 132 | 329 | - | - | - |
| Number of small ruminants | 0 | 132 | 329 | - | - | - |
|  | 1-5 | 58 | 89 | 2.65 | 0.274 | 1.52 - 4.61 |
|  | >5 | 27 | 16 | 14.83 | 0.001 | 4.86 – 45.23 |
| Have aborted animals in last 12 months | Yes | 51 | 42 | 9.65 | 0.001 | 3.37 – 27.64 |
|  | No | 55 | 118 | - | - | - |
| Number of aborted animals in last 12 months | 0 | 55 | 118 | - | - | - |
|  | 1 | 26 | 39 | 6.17 | 0.715 | 2.06 – 18.48 |
|  | >1 | 25 | 3 | 52.70 | 0.001 | 11.32– 245.4 |
| Notification of authorities of a case of abortion | Yes | 7 | 28 | 0.14 | 0.002 | 0.04 – 0.45 |
|  | No | 44 | 27 | - | - | - |
| Have infected animals with *Brucella* spp. in last 12 months | Yes | 6 | 0 | >99.9 | 0.98 | <0.01- >99.9 |
|  | No | 100 | 160 | - | - | - |
| Notification of authorities of a case of *Brucella* spp. | Yes | 8 | 46 | <0.001 | 0.99 | <0.0- >99.9 |
|  | No | 2 | 0 | - | - | - |
| Protective measures for animals against *Brucella* spp. infection | Yes | 38 | 111 | 0.11 | 0.0001 | 0.04 - 0.31 |
|  | No | 68 | 49 | - | - | - |
| Vaccinate animals against brucellosis | Yes | 19 | 93 | 0.1 | 0.0001 | 0.04- 0.24 |
|  | No | 87 | 67 | - | - | - |
| Involve in activities where you come in contact with animals | Yes | 136 | 148 | 6.85 | 0.0001 | 4.13 – 11.35 |
|  | No | 81 | 286 | - | - | - |
| protective measures applied if come in contact with animals | Yes | 119 | 127 | 6.15 | 0.0001 | 3.70 – 10.25 |
|  | No | 98 | 307 | - | - | - |
| number of protective measures you apply when come in contact with animals | 0 | 98 | 307 | - | - | - |
|  | 1-2 | 23 | 71 | 1.23 | 0.008 | 0.59 – 2.57 |
|  | >2 | 96 | 56 | 20.08 | 0.0001 | 9.03 – 44.64 |
| participating in dairy product processing | Yes | 76 | 141 | 4.06 | 0.037 | 1.09– 15.08 |
|  | No | 141 | 293 | - | - | - |
| have consumed dairy products over the last 3 months | Yes | 210 | 378 | 5.28 | 0.002 | 2.12 – 12.62 |
|  | No | 7 | 56 | - | - | - |
| score of milk consumption | 0 | 9 | 58 | - | - | - |
|  | 1-2 | 75 | 234 | 2.52 | 0.18 | 1.15 – 5.54 |
|  | >2 | 133 | 142 | 12.49 | 0.0001 | 5.28 – 29.57 |
| consumed unpasteurized milk | Yes | 21 | 7 | 6.17 | 0.0001 | 2.62 – 14.55 |
|  | No | 196 | 427 | - | - | - |
| have consumed fermented milk | Yes | 163 | 279 | 1.7 | 0.006 | 1.15 – 2.4 |
|  | No | 54 | 155 | - | - | - |
| have consumed homemade cheese | Yes | 141 | 189 | 3.61 | 0.0001 | 2.33 – 5.67 |
|  | No | 76 | 245 | - | - | - |
| have consumed yoghurt | Yes | 63 | 71 | 4.32 | 0.0001 | 2.3 – 8.13 |
|  | No | 154 | 363 | - | - | - |
| have consumed cream | Yes | 84 | 131 | 1.44 | 0.035 | 1.03 – 2.01 |
|  | No | 133 | 303 | - | - | - |
| Protective measures take to protect humans from infection | Yes | 92 | 153 | 1.53 | 0.037 | 1.03 – 2.30 |
|  | No | 125 | 281 | - | - | - |
| Number Protective measures take to protect humans from infection | 0 | 125 | 281 | - | - | - |
|  | 1-2 | 83 | 104 | 2.30 | 0.0001 | 1.46 – 3.85 |
|  | >2 | 9 | 49 | 0.35 | 0.0004 | 0.15 – 0.79 |
| Infected household in last 3 months | Yes | 10 | 8 | 2.68 | 0.048 | 1.01 – 7.09 |
|  | No | 207 | 426 | - | - | - |

The risk of having human brucellosis infection was 9.65 times higher among participants reporting the occurrence of animal abortion [(95% CI: 3.37 – 27.64); *p* < 0.001]. This risk mounted to 52.7 times more among cases who have high number of aborted animals comparing to those having utmost a single aborted animal [95% CI: 11.32 – 245.4); *p* < 0.001]. On the other hand, people who have vaccinated animals [OR (95% CI) = 0.1 (0.04 – 0.24)], follow protective measures for animals against *Brucella* spp. infection [OR (95% CI)= 0.11 (0.04 – 0.31)] and notify authorities of in case of animal abortion [OR (95% CI)= 0.14 (0.04 – 0.45)] were significantly at lower risk of getting human brucellosis infection (*p* < 0.05) (Table 6). Nevertheless, these 5 variables; have aborted animals, number of aborted animals, follow protective measures with animals, vaccinate animals against brucellosis and notify authorities of case of abortion were not incorporated in the final multivariable analysis because in each variable, 385 out of 651 respondents had no animals in their households.

The presence of animals in the households of respondent, number of owned animals, presence of small ruminants and number of small ruminants were found to have strong collinearity at *p* < 0.05. In the final multivariate logistic regression model, these variables were detected as confounders to each other. Thus only “the presence of small ruminants in respondent household” was included in the final model as it was the only variable with constant OR (95% CI) and *p* value comparing to other variables. Likewise, strong collinearity was found between the use of protective measures and number of protective measures followed by the livestock owners when come in contact with animals, with the first variable being used in the final multivariate analysis. On the other hand, the number of protective measures that a person take to prevent human infection was considered in the final model instead of if the person follows these protective measures or not for the same last reason. Milk score consumption was removed from the multivariate analysis for the same reason as before since they had significantly collinearity but act as confounders for the variables: consumption of dairy products in last 3 months, consumption of unpasteurized milk, consumption of yoghurt and consumption of home-made cheeses. Likewise, having infected animal in the households, and notification of authorities for of a case of Brucella were found not significantly accounted as risk factors for human infection with brucellosis and were removed from the multivariate analysis as their *p* value was > 0.2. However, we do not think that having infected animal is not considered a risk factor for brucellosis infection but the results we had may be due to the few number of respondents to this question.

The following variables were used in the multivariate model: presence of small ruminants in the households, if the participant involved in activities where he or she comes in contact with animals, if the respondent follows protective measures when he/she come in contact with animals, number of protective measures the respondent apply to protect himself and household members from *Brucella* infection, if the participant used to consume dairy products in the last 3 months before the test, if the participant normally consume unpasteurized milk, fermented milk, cream, home-made chees and yoghurt, if the participant involve in milk products processing and if the participant has an infected household member with *Brucella* spp. in last 3 months before the meeting. Involvement in contact with animals was found as confounding factor for presence of small ruminant, so the latter variable was removed. Testing for confounder was carried out by monitoring the change of logit of factors by removing a suspected factor from the model. Out of the 11 variables, 5 were removed before building the final model because they did not meet the 0.05 significance level for consideration into the final model; consumption of fermented milk and cream, involvement in dairy products processing, infected household in last 3 months, and protective measures followed when the participant come in contact with animals.

Results of multivariate logistic regression analysis are shown in (Table 7). Consumption of dairy products in the last 3 months before the test and consumption of unpasteurized milk or home-made cheeses and yoghurt were associated with higher odds of having brucellosis infection [OR (95% CI)= 2.71 (1.06-6.93); *p* < 0.038] and [OR (95% CI)= 4.12 (1.62 - 10.75); *p*< 0.003, 1.96 (1.17- 3.30); *p* < 0.011 and 2.51 (1.21 – 5.24); *p* < 0.014] respectively. Participants who were involved in activities where they come in contact with animals had more than 4.97 times greater odds of having brucellosis infection [(95% CI)= (2.84 - 8.72); *p* < 0.001]. Finally, participants who take more protective actions for themselves against brucellosis are almost 5 times less likely to have been diagnosed with brucellosis [OR (95% CI)= 0.23 (0.10 - 0.58); *p* < 0.001].

**Table (7):**
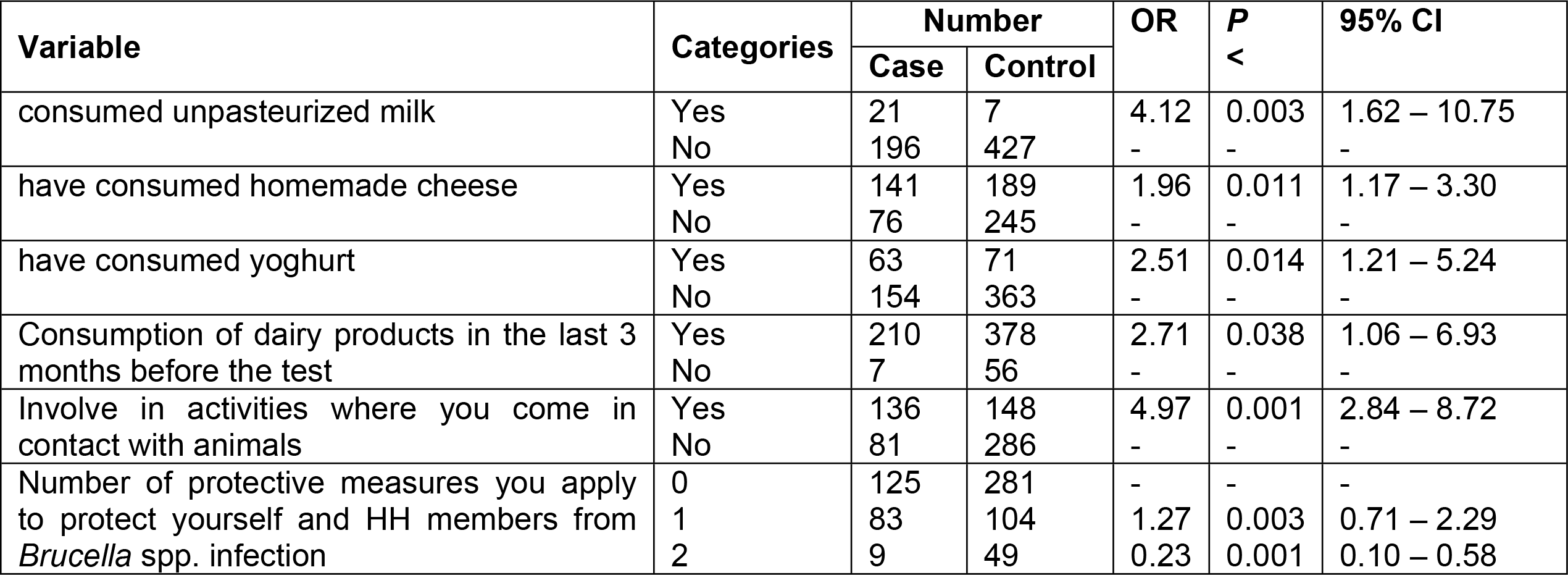
Results of multivariable model for the association between selected potential risk factors and individual Brucella spp. positive status

We developed another model to test the association between selected potential risk factors and individual *Brucella* spp. positive status among people who owned animals at their households. We used in such model the same variables including these 5 variables deleted from the first model (Table 8). In addition to the results of previous model results, in the second model, vaccination of animals and if the owner has aborted animals and notification of abortion were removed from the final model as they were confounders for the variables: protective measures respondents apply to prevent their animals from *Brucella* spp. infection and the number of abortion per animal, respectively. Increase in the number of abortions per animal increased the chance for brucellosis infection among human cases by 49.33-fold [95% CI= (1.5– 155.7); *p* < 0.0001] while the practice of protective measures with animals was protective [OR (95% CI) = 0.11 (0.03-0.45); *p* < 0.002].

**Table (8):** Results of multivariable model for the association between selected potential risk factors and individual Brucella spp. seropositive status among people who owned animals

| Variable | Categories | Number |  | OR | P<br>< | 95% CI |
| --- | --- | --- | --- | --- | --- | --- |
|  |  | Case | Control |  |  |  |
| Number of abortions per animal | 2 | 25 | 3 | 49.33 | 0.001 | 8.79 – 276.91 |
|  | 1 | 26 | 39 | 4.80 | 0.46 | 1.5 – 15.57 |
|  | 0 | 55 | 118 | - | - | - |
| protective measures you apply to protect animals from *Brucella* spp. infection | Yes | 38 | 111 | 0.11 | 0.002 | 0.03 – 0.45 |
|  | No | 68 | 48 | - | - | - |

## Discussion

Despite its high burden in many parts of the world, brucellosis is rarely prioritized by health systems in developing countries and is considered a neglected zoonosis [18,19]. The current study adopted an integrated approach in attempt to address the gap in knowledge, attitude and practice towards brucellosis from both veterinary and human health stand point in the Nile Delta in Egypt, where the disease remains endemic [8–13,15,16,20–23]. In this major agrarian region, most of residents rely on agriculture with a larger proportion entirely depending on livestock production for their livelihood and these animals could pose a public health threat to humans. The present study revealed that 40.7% of the study population keep animals in their households and they accommodate cows and buffaloes with sheep and goats. This represents a great risk factor for brucellosis transmission between animals and to humans since small ruminants are the primary hosts of B. *melitensis*, the predominant *Brucella* spp. circulating in Egypt and cattle are more likely spillover hosts [10]. Moreover, villagers use mobile village flocks of small ruminants for breeding with household animals thus maintaining high level of Brucella in these animals [10].

Knowledge about the disease and preventive herd management practices have previously been identified as the most important factors required for minimizing the disease risk in animals [24] Apparently, brucellosis as a disease of animals is not known by our study population, since more than two thirds of the participants had not heard of the disease or its possible transmission. This is consistent with previous reports in Egypt[25] and Nigeria[26,27] but differed from others studies conducted in Kenya[28], Jordon[29], Egypt[10,12], India[30], Tajakistan[31] and Nigeria[27] where the majority of the study respondents had heard of brucellosis and correctly believed that brucellosis is transmissible from animals to humans. This could be attributed to different population composition since these studies tended to interview pastoralist, shepherds and livestock keepers whereas more than half of our study population had no contact with animals.

Poor knowledge regarding the etiology and transmission of brucellosis could negatively impact on individuals’ preventive and control methods of the disease at the humans-animal interface due to misconception on its determinants. This was evident in the current study since participants’ knowledge significantly reflected their attitude, practice, perceived barriers and perceived susceptibility and risk. However, high levels of awareness not necessarily go hand in hand with accurate behavior and practices, as the perception of a risk is influenced by many factors such as life experience and culture [32]. Contrary to findings in Tajikistan[31] and Yemen[33], participants awareness about the disease did not correlate with the educational level, although participants with a low level of literacy were more likely to have risky practices and high perceived barriers towards disease control. Those with a lower level of education are thus likely at higher risk of contracting brucellosis.

Similar to reports from Uganda[34] and Tajikistan[31], the main sources of information on brucellosis in the present study area was neighbors and veterinarians. Few participants in the current study mentioned media, such as radio or television as a source of information about the disease, although they were stated as the main source concerned with disseminating information on brucellosis disease in Jordan [35]. It is worth noting that in Egypt-from our experience as Egyptian authors-the media typically disseminate health information only in cases of epidemics, but diseases of endemic nature are always neglected. Together, these highlight the powerful role of veterinarians or the community health workers play in terms of relaying important health messages to livestock owners in the study area who in most circumstances have challenges in accessing basic health care services particularly the cost. Deliberate actions should therefore be undertaken to incorporate all aspects of health care education for the livestock owners.

Furthermore, livestock keepers in Egypt do not call veterinarian in case of abortion or parturition. Instead, they and others in the village assist with calving, usually by pulling the calf out or removing placenta and foetal membranes. Most farmers dispose placentas, aborted fetuses and carcasses in the water canals or bury them. Brucella species can survive in aborted fetuses and humid environment (manure and soil) for a period up to 8 months [36]. Most villagers come in contact with this potentially contaminated water through daily routines such as bathing, irrigation of fields, washing of utensils, fishing and other activities. Therefore, lack of effective carcass disposal and unrestricted local husbandry methods could result in significant environmental contamination with *brucella* and increases the risk of disease transmission to human and livestock populations [37].

Villagers never wear protective gloves or masks when assisting with the parturition or abortion of animals or whilst handling placentas and aborted fetuses [10]. Interestingly, cases were more likely to admit the use of protective measures including gloves and mask and emphasized washing their hands more often comparing to controls suggesting a reverse causality although our proposed model revealed that those who practice 2 or more protective measures were at lower risk of getting brucellosis. Moreover, human brucellosis cases could have experienced a major illness and thus now protect themselves. This could otherwise be explained by the “Hawthorne effect,’’ that is, “a behavioral tendency of subjects to provide information consistent with their perception of the study objectives that positively value hygienic behaviors” [38]. The deficient use of protective equipment is attributed not only to poor knowledge of the risk with this practice but also the lack of access to protective clothing like gloves.

Knowledge of the animal species affected and signs or symptoms of brucellosis in animals are crucial because it positively impacts on livestock owners’ practices towards prevention and control measures of brucellosis in both animals and humans. In this regard, the basic knowledge of interviewed participants about the animal species that could be affected by brucellosis was poor. This finding contrasts with the findings of studies in Tajikistan[31] and an earlier study in Egypt [10] where the majority of respondents knew that cattle, sheep and goats could be affected. Few mentioned fish as a susceptible host to brucellosis. Although this may appear incorrect, Nile catfish have been found to be infected with B. *melitensis* in small tributaries of Nile canals in Kafrelsheikh, Gharbiya, Menoufia, and Dakahlia governorates in the Nile Delta region. It was isolated from liver, kidney, spleen samples and skin swabs of wild fish; but not from samples of farmed fish [37]. This indicates that the heavy contamination of water by animal waste presents a new potential route of human infection. Other respondents stated cats, dogs and rats as susceptible host for brucellosis. In fact, Brucella *melitensis* biovar 3 has previously been isolated from stray dogs and rats trapped near dairy farms and water canals in Egypt; at levels higher than seropositive herds [39]. No doubt that responded stated these rare brucellosis susceptible species are not acquainted with what have been recently published in the literature and certainly reported that by chance. However, this implies the need for increasing the knowledge and awareness of the community regarding these emerging issues.

Kozukeev et al., (2006) found that good knowledge of mode of transmission of brucellosis from animals to humans had a protective effect towards human infection [40]. In our study, among those who were aware of the zoonotic nature of brucellosis, consumption of raw contaminated milk and milk products, contact with infected ruminants and involvement in infected animal abortion or parturition were the most frequently listed modes of brucellosis transmission. The participants’ response regarding involvement in animal husbandry and consumption of milk as a mode of transmission was comparable to earlier findings in Egypt[10,12], Kenya[28] and Uganda[34,41]. Other additional routes were mentioned most of which have been identified in many studies as major risk factors for transmission of brucellosis at human-animal interface [5,33,40,42–45].

Unregulated buying and selling of animals and animal products are great hazards as they facilitate transmission between new animals and to people [10,46]. Regarding farmers practice when the animals abort or become infected, in the current study, a considerable number of livestock owners sold the aborted or infected animal for reproduction for other livestock owners or for slaughtering for butchers. This imposes a great risk of infection among abattoir workers and butchers. These findings were in the same line with Holt et al., [10] in Egypt. Animals purchased at a market can move without restriction to anywhere in Egypt. This may increase the transmission of brucellosis, not only between households in the same village, but also between villages and even larger geographical areas [25].

Only a small proportion of the study respondents perceived that brucellosis was a serious disease in both animals and humans and that animal husbandry is a risky practice. Accordingly, they had unfavorable attitude towards good practices in prevention of brucellosis. Several known high-risk behaviors were common self-reported practices among the study participants, particularly cases. They were more likely to engage in risky practices that could expose them to infection. This was evident from actions most of them would take when confronted with an aborting animal in their herd. The majority would not seek governmental veterinary services and thus would not notify the disease. Some manage animal infection or abortion on their own and some call private veterinarians. Holt et al., (2011) stated that private veterinarians do not be report brucellosis as they are unlikely to be penalized for that and sometime get benefits from livestock owners for not reporting. In our experience, livestock owners fear of economic losses caused by governmental tracing and culling of their livestock. Private veterinarians usually reside within the village they work in and know the livestock owners, who is often a friend or even a relative and thus have loyalty for them. They advise livestock owners to fatten suspected animals to be able to sell them for slaughtering but not to get the animal tested thus facilitating the underreporting and disease surveillance [10]. Livestock owners find it is easier and more profitable to sell animals than to notify the veterinary authorities and wait until they test and slaughter the positive animal. In fact, test and slaughter strategy implemented in Egypt guarantees compensation for livestock owner. However, it accounts for less than 50% of the market value of the animal and often takes long time to receive [10,47]. This results in underreporting of the diseases and hence hinders brucellosis control in Egypt. The same issue is reported in Greece where 44% of patients with brucellosis would not allow veterinary investigation as they were worried about the effects on their herd [48]. The government should approach private veterinarians and work more closely with them in order to improve the flow of information and disease notification. Furthermore, adequate compensation or replacement animals should be considered [10].

Female animals infected with *Brucella* spp. excrete high concentrations of the organism in their milk, placental membranes and aborted fetuses [19,49]. Therefore, there is a risk of humans becoming infected through consumption of dairy products. In fact, consumption of raw milk has been previously described as one the riskiest practices [50]. In the present study, drinking raw fresh milk was uncommon practice owing to the awareness of its hazards. Nevertheless, some traditionally have the concept that consuming raw milk is healthier, boosts immunity and have a cooling effect in the summer. Consumption of unpasteurized milk was practiced by more cases comparing to controls. This risk appeared negligible in an earlier Egyptian study [12]. However, in Kenya, nearly all respondents consumed raw milk [28]. More education of villagers is needed to protect themselves from the exposure and to reduce the risk of facilitating the transmission and spread of brucellosis.

Consistent with findings from Tajikistan [31], the majority of households admitted their involvement in dairy product processing and sold unpasteurized dairy products from farms directly to consumers on regular basis. Commercialization of these homemade products is not restricted to the nearby villages but is remotely marketed without surveillance and consequently without proper refrigeration, conservation, packaging or storage. Raw milk and milk products are surplus for sale in the Egyptian market and are of a high demand for home consumption in big urban cities including Cairo and Alexandria. Consumers just heat or boil raw milk before its consumption. Such system contributes around 72% of total milk produced in Egypt. Even though the legislation in Egypt imposes the pasteurization of milk before processing, only modern large-scale dairy plants and 27% of the municipal dairy plants follow the law instructions [51]. This practice of trading with unpasteurized milk and home-made animal products could constitute a risk to public health and put food safety at risk.

Although having infected animal in the households, and notification of authorities for of a case of brucella were removed from the logistic regression model for predicting brucellosis infection, we do not think that these are not considered as risk factors for brucellosis infection but the results we had may be due to the fewer number of respondents to these questions.

The number of abortions per animal increased the chance for infection among human cases. This is an important finding in the present work especially for people who insist to keep aborted animals [for fattening or reproduction] and do not like to slaughter them. It is worth noting that livestock owners deny having animal abortion to be able to sell them or their milk [12].

Animal vaccination appeared as a protective factor in our proposed logistic regression model for risk factors of human infection. This finding supports the economic and health benefits of animal vaccination in reducing the occurrence of human brucellosis [52]. Importantly, a significant number among those actually practicing animal vaccination did not list this practice among the measures that they adopt for protecting themselves, their household member or their animals against brucellosis. This means that villagers can follow a preventive measure without being aware of its benefits probably because this was not properly explained to them.

The key limitation of the present work is the self-reporting on practices by the respondents that was subject to recall bias, Hawthorne effect and the face-to-face interview situation. Observational checklist could have enhanced this type of bias in assessing attitudes and behaviors.

## Conclusion and Perspectives

Findings from this study demonstrate a poor understanding of brucellosis and a high level of risky practices being undertaken on farms and at households across a wide region of the country. These all conflict with disease intervention strategies and contribute to the risk of humans to contract brucellosis. Lack of compliance of the villagers and small livestock keepers with the disease control measures stands behind the lack of success of the current national control program for brucellosis in Egypt. Particularly, underreporting animal infection and abortion; underutilization of animal vaccination service; unsanitary disposal of abortus; use of milk of infected/aborted ruminants and lack of protective measure when practicing animal husbandry. Understanding of the knowledge, attitudes and practices is crucial for assessing the feasibility, acceptability and barriers of potential measures that might be instituted. This strongly supports the need for including health education as part of brucellosis control programs in rural communities with a special emphasis on hygienic animal husbandry, disease notification and the benefits of human animal vaccination. This can be achieved by targeted messages in local FM radios and television, besides integrating the community health volunteers in the control and prevention efforts.

The prospective of this work is to collect information from different community sections to assess the economic impacts of brucellosis on small livestock holders and on the national level. Data of the current study will be the corner stone for the following section of the project which aims at building a more realistic to apply and economically efficient socioeconomic model for brucellosis control in the Nile Delta region and consequently in Egypt to substitute the current national brucellosis control strategy.

## Ethical considerations

### Compliance with Ethical Standards

The study was approved by the institutional review board and the Ethics Committee of the High Institute of Public Health-Alexandria University [no. 14-2014/6]. Permission to conduct the study was obtained from the Ministry of Health. The research was conducted in accordance with the ethical guidelines of the Declaration of Helsinki (2013) and the International Conference on Harmonization Guidelines for Good Clinical Practice. All participants were given verbal information about the study and were assured about the confidentiality, protection and anonymity of their data. Informed written consent was obtained from the individuals participating voluntarily. Data sheets were coded to ensure anonymity and confidentiality of patient’s data.

This article does not contain any studies with animals performed by any of the authors.

### Data availability

All data are fully available without restriction by the corresponding author at and through the public data repository Harvard Dataverse at https://dataverse.harvard.edu/dataverse/BrucellosisEgypt

## Acknowledgements

We would like to acknowledge the study participants, the Ministry of Health and Population, and the District Veterinary office for the support provided during the investigations.

## Supporting Information Legends

S1 File: questionnaire tool developed by the authors for data collection

S2 Checklist: STROBE Checklist

## Supplementary tables (for online display)

**Table S1:** Sociodemographic characteristics of household members of the study population

|  | Total<br>(n=651) |  | Cases<br>(n=217) |  | Controls<br>(n=434) |  | *\*p-value* |
| --- | --- | --- | --- | --- | --- | --- | --- |
|  | No. | % | No. | % | No. | % |  |
| Number of household member exposed to *Brucella* in last 3 months |  |  |  |  |  |  |  |
| Yes | 18 | 2.8 | 10 | 4.6 | 8 | 1.8 | 0.015 |
| No | 588 | 90.3 | 186 | 85.7 | 402 | 92.6 |  |
| Do not know | 45 | 6.9 | 21 | 9.7 | 24 | 5.5 |  |
| household member infected with *Brucella* in last 3 months (n=18) |  |  |  |  |  |  |  |
| Mother | 3 | 16.7 | 2 | 20.0 | 1 | 12.5 | 0.001 |
| Sister | 2 | 11.1 | 2 | 20.0 | 0 | 0.0 |  |
| Brother | 4 | 22.2 | 4 | 40.0 | 0 | 0.0 |  |
| Cousin | 7 | 38.9 | 1 | 10.0 | 6 | 75.0 |  |
| Son | 2 | 11.1 | 1 | 10.0 | 1 | 12.5 |  |
| Age (n=18) |  |  |  |  |  |  |  |
| 12-<25 | 6 | 33.3 | 3 | 30.0 | 3 | 37.5 | 0.94 |
| 25-<40 | 7 | 38.9 | 4 | 40.0 | 3 | 37.5 |  |
| 40--55 | 5 | 27.8 | 3 | 30.0 | 2 | 25.0 |  |
| Gender (n=18) |  |  |  |  |  |  |  |
| Male | 12 | 66.7 | 6 | 60.0 | 6 | 75.0 | 0.45 |
| Female | 6 | 33.3 | 4 | 40.0 | 2 | 25.0 |  |
| Education (n=18) |  |  |  |  |  |  |  |
| Illiterate | 3 | 16.7 | 3 | 30.0 | 0 | 0.0 | 0.007 |
| Primary | 2 | 11.1 | 1 | 10.0 | 1 | 12.5 |  |
| Secondary | 3 | 16.7 | 3 | 30.0 | 0 | 0.0 |  |
| University | 10 | 55.6 | 3 | 30.0 | 7 | 87.5 |  |
| Occupation (n=18) |  |  |  |  |  |  |  |
| Not currently employed | 1 | 5.6 | 0 | 0.0 | 1 | 12.5 | 0.085 |
| student | 3 | 16.7 | 1 | 10.0 | 2 | 25.0 |  |
| housewife | 4 | 22.2 | 4 | 40.0 | 0 | 0.0 |  |
| farmer | 3 | 16.7 | 2 | 20.0 | 1 | 12.5 |  |
| employee | 4 | 22.2 | 2 | 20.0 | 2 | 25.0 |  |
| professional | 1 | 5.6 | 0 | 0.0 | 1 | 12.5 |  |
| clerical work | 2 | 11.1 | 1 | 10.0 | 1 | 12.5 |  |

*\*p*-value is for chi square test

**Table S2:**
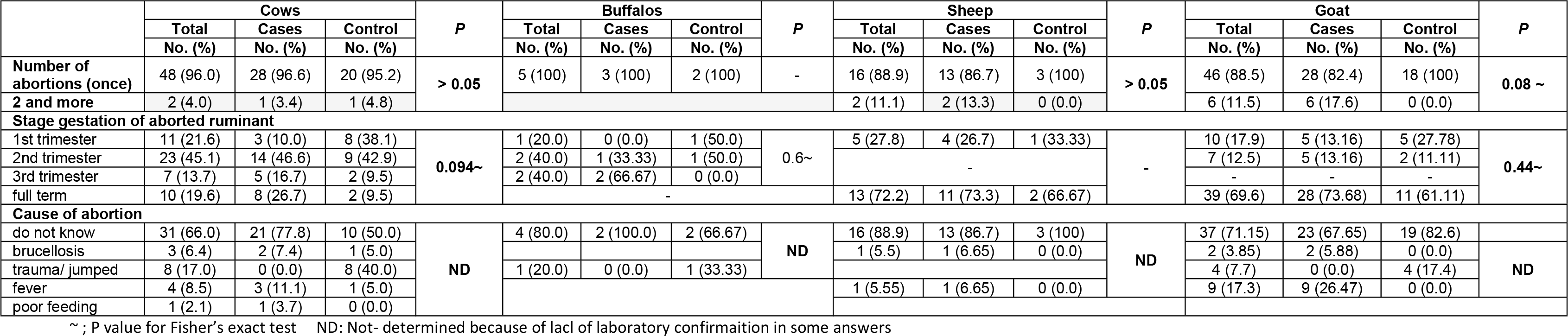
Occurrence of abortion among animals owned by the study participants

**Table S3:**
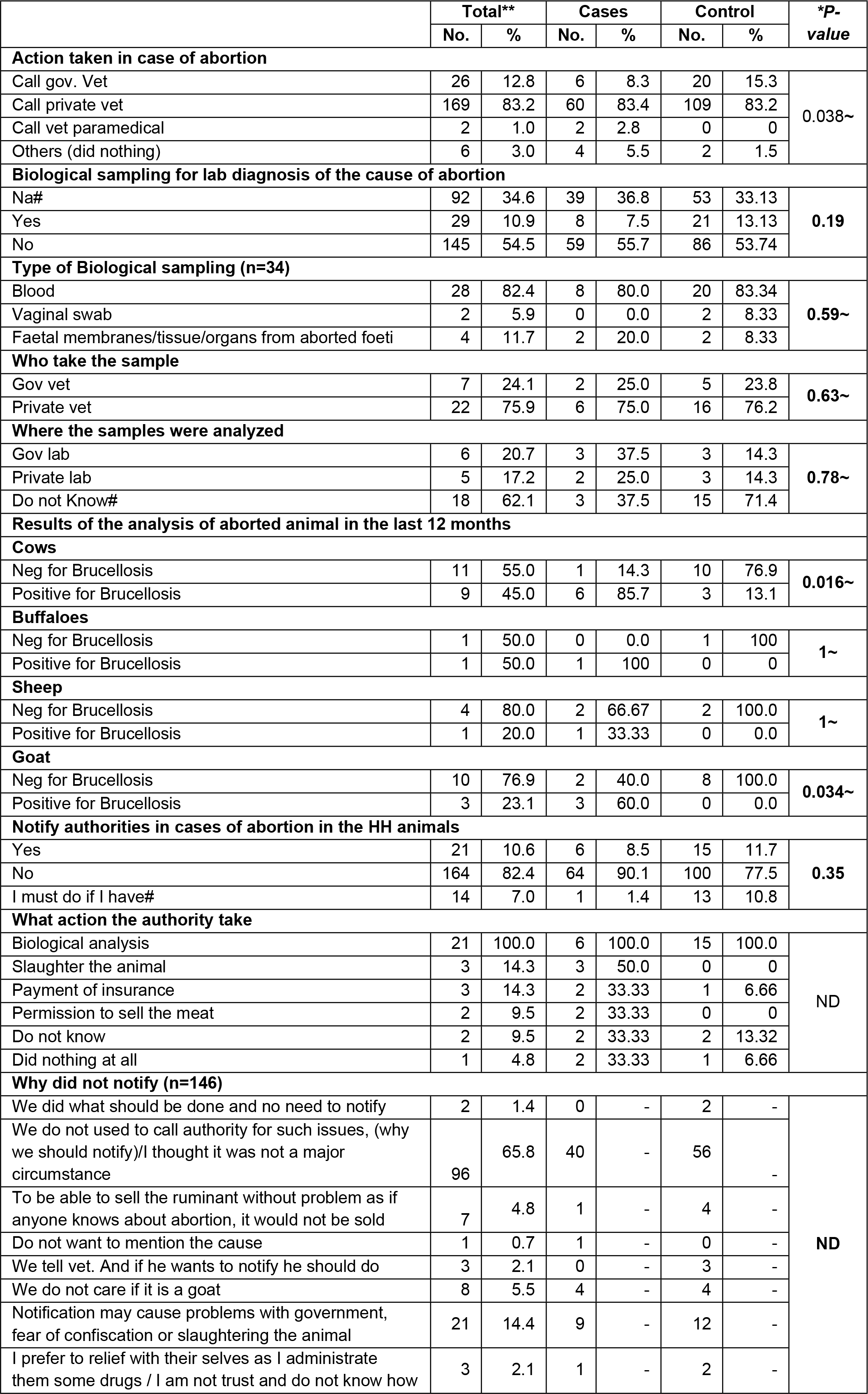
Actions taken by cases and controls in case of animal abortion

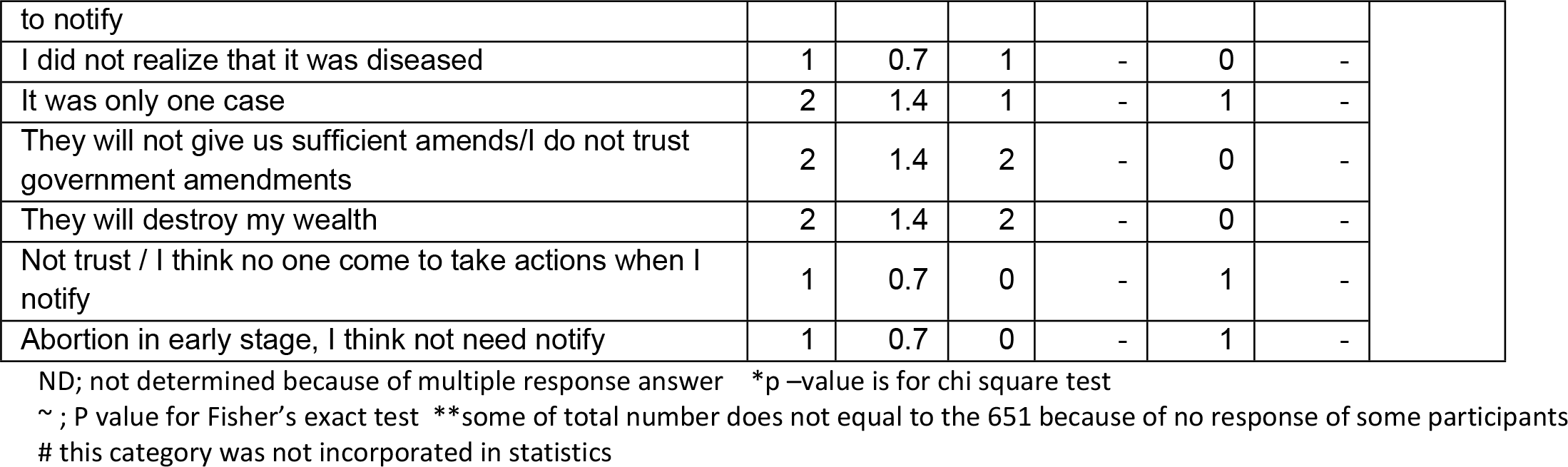

**Table S4:**
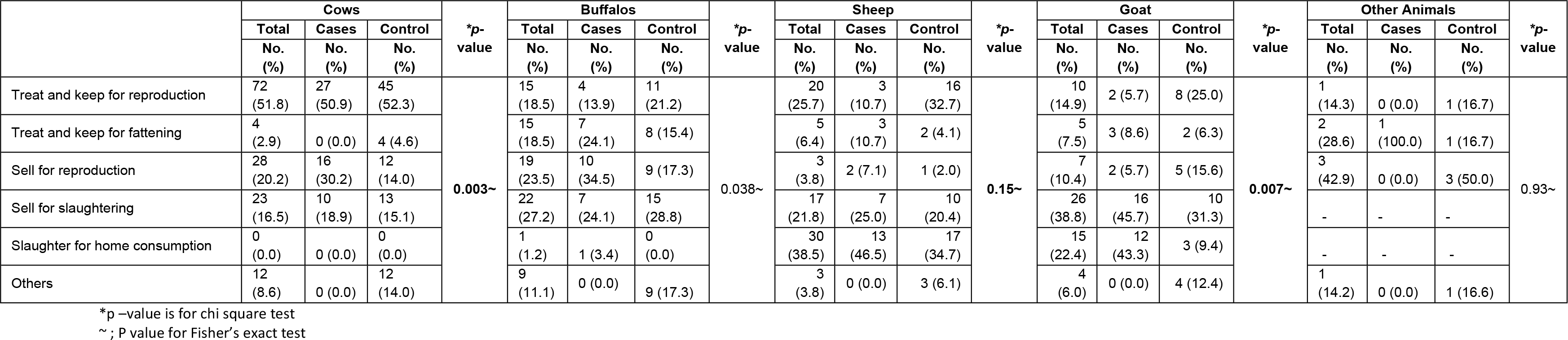
What cases and control do with aborted animals

**Table S5:**
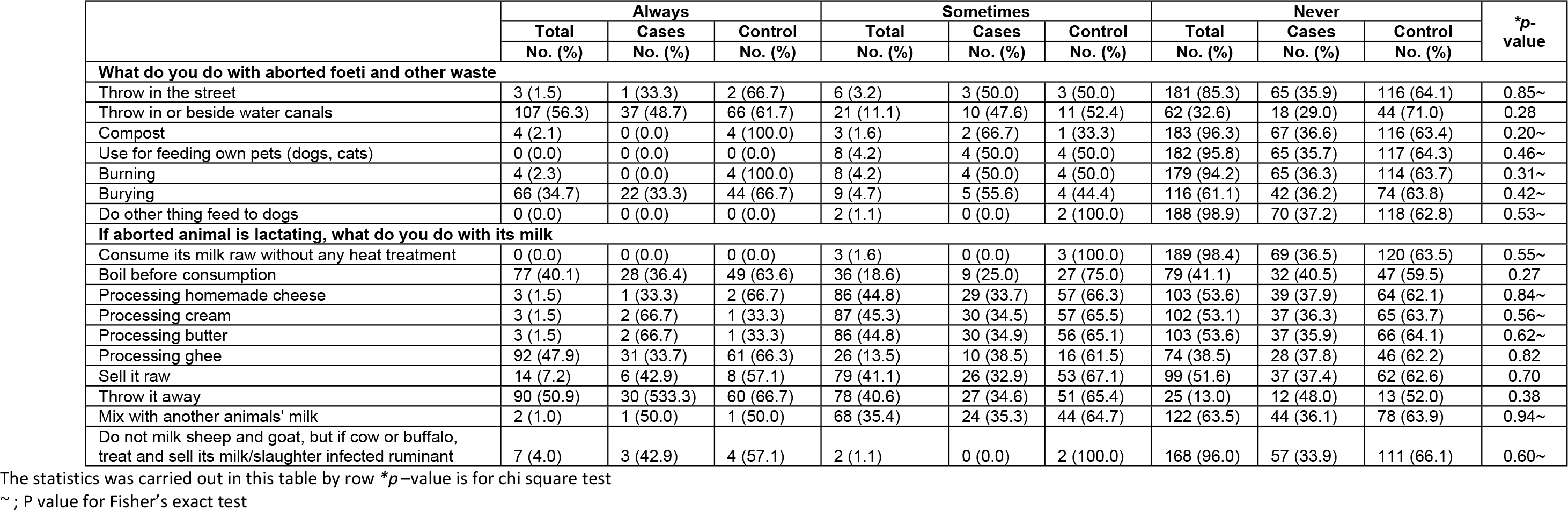
How cases and controls handle aborted foeti, other waste and milk from aborted animals

**Table S6:**
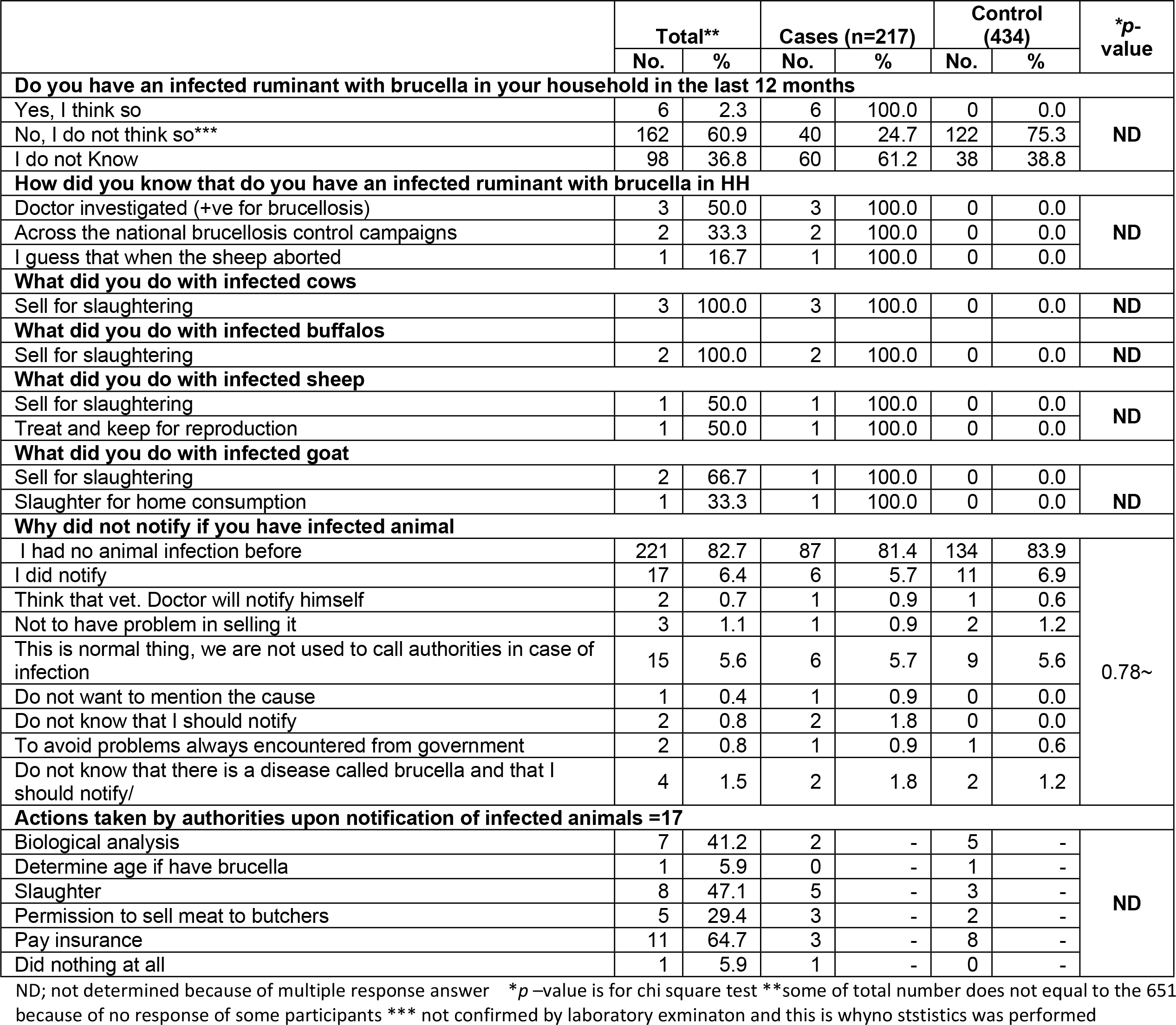
Occurrence of animal brucellosis in households of cases and controls

|  | Total** |  | Cases (n=217) |  | Control (434) |  | *p-value |
| --- | --- | --- | --- | --- | --- | --- | --- |
|  | No. | % | No. | % | No. | % |  |
| **Do you have an infected ruminant with brucella in your household in the last 12 months** |  |  |  |  |  |  |  |
| Yes, I think so | 6 | 2.3 | 6 | 100.0 | 0 | 0.0 | ND |
| No, I do not think so*** | 162 | 60.9 | 40 | 24.7 | 122 | 75.3 |  |
| I do not Know | 98 | 36.8 | 60 | 61.2 | 38 | 38.8 |  |
| **How did you know that do you have an infected ruminant with brucella in HH** |  |  |  |  |  |  |  |
| Doctor investigated (+ve for brucellosis) | 3 | 50.0 | 3 | 100.0 | 0 | 0.0 | ND |
| Across the national brucellosis control campaigns | 2 | 33.3 | 2 | 100.0 | 0 | 0.0 |  |
| I guess that when the sheep aborted | 1 | 16.7 | 1 | 100.0 | 0 | 0.0 |  |
| **What did you do with infected cows** |  |  |  |  |  |  |  |
| Sell for slaughtering | 3 | 100.0 | 3 | 100.0 | 0 | 0.0 | ND |
| **What did you do with infected buffalos** |  |  |  |  |  |  |  |
| Sell for slaughtering | 2 | 100.0 | 2 | 100.0 | 0 | 0.0 | ND |
| **What did you do with infected sheep** |  |  |  |  |  |  |  |
| Sell for slaughtering | 1 | 50.0 | 1 | 100.0 | 0 | 0.0 | ND |
| Treat and keep for reproduction | 1 | 50.0 | 1 | 100.0 | 0 | 0.0 |  |
| **What did you do with infected goat** |  |  |  |  |  |  |  |
| Sell for slaughtering | 2 | 66.7 | 1 | 100.0 | 0 | 0.0 | ND |
| Slaughter for home consumption | 1 | 33.3 | 1 | 100.0 | 0 | 0.0 |  |
| **Why did not notify if you have infected animal** |  |  |  |  |  |  |  |
| I had no animal infection before | 221 | 82.7 | 87 | 81.4 | 134 | 83.9 | 0.78~ |
| I did notify | 17 | 6.4 | 6 | 5.7 | 11 | 6.9 |  |
| Think that vet. Doctor will notify himself | 2 | 0.7 | 1 | 0.9 | 1 | 0.6 |  |
| Not to have problem in selling it | 3 | 1.1 | 1 | 0.9 | 2 | 1.2 |  |
| This is normal thing, we are not used to call authorities in case of infection | 15 | 5.6 | 6 | 5.7 | 9 | 5.6 |  |
| Do not want to mention the cause | 1 | 0.4 | 1 | 0.9 | 0 | 0.0 |  |
| Do not know that I should notify | 2 | 0.8 | 2 | 1.8 | 0 | 0.0 |  |
| To avoid problems always encountered from government | 2 | 0.8 | 1 | 0.9 | 1 | 0.6 |  |
| Do not know that there is a disease called brucella and that I should notify/ | 4 | 1.5 | 2 | 1.8 | 2 | 1.2 |  |
| **Actions taken by authorities upon notification of infected animals =17** |  |  |  |  |  |  |  |
| Biological analysis | 7 | 41.2 | 2 | - | 5 | - | ND |
| Determine age if have brucella | 1 | 5.9 | 0 | - | 1 | - |  |
| Slaughter | 8 | 47.1 | 5 | - | 3 | - |  |
| Permission to sell meat to butchers | 5 | 29.4 | 3 | - | 2 | - |  |
| Pay insurance | 11 | 64.7 | 3 | - | 8 | - |  |
| Did nothing at all | 1 | 5.9 | 1 | - | 0 | - |  |

ND; not determined because of multiple response answer \*p –value is for chi square test \*\*some of total number does not equal to the 651 because of no response of some participants \*\*\* not confirmed by laboratory examination and this is why no statistics was performed

**Table S7:** How cases and controls handle milk from infected ruminants

|  | Always |  |  | Sometimes |  |  | Never |  |  | *p-value |
| --- | --- | --- | --- | --- | --- | --- | --- | --- | --- | --- |
|  | Total | Cases | Control | Total | Cases | Control | Total | Cases | Control |  |
|  | No. (%) | No. (%) | No. (%) | No. (%) | No. (%) | No. (%) | No. (%) | No. (%) | No. (%) |  |
| if infected animal is lactating, what do you do with its milk |  |  |  |  |  |  |  |  |  |  |
| consume its milk raw without any heat treatment | 0 (0.0) | 0 (0.0) | 0 (0.0) | 4 (2.2) | 0 (0.0) | 4 (100.0) | 173 (97.7) | 60 (34.7) | 113 (65.3) | 0.30~ |
| boil before consumption | 74 (41.6) | 26 (35.1) | 48 (64.9) | 27 (15.2) | 6 (22.2) | 21 (77.8) | 77 (43.3) | 29 (37.7) | 48 (62.3) | 0.33 |
| processing homemade cheese | 1 (0.6) | 1 (100.0) | 0 (0.0) | 82 (44.6) | 27 (32.9) | 55 (67.1) | 95 (53.3) | 33 (34.7) | 62 (65.3) | 0.47~ |
| processing cream | 1 (0.6) | 1 (100.0) | 0 (0.0) | 83 (45.1) | 27 (32.5) | 56 (67.5) | 94 (52.8) | 33 (35.1) | 61 (64.9) | 0.39~ |
| processing ghee | 2 (1.1) | 1 (50.0) | 1 (50.0) | 24 (13.4) | 8 (33.3) | 16 (66.7) | 152 (85.) | 52 (34.2) | 100 (65.8) | 0.57~ |
| processing butter | 1 (0.6) | 1 (100.0) | 0 (0.0) | 84 (47.2) | 28 (33.3) | 56 (66.7) | 93 (52.2) | 32 (34.0) | 61 (65.6) | 0.52~ |
| sell it raw | 13 (7.3) | 5 (38.5) | 8 (61.5) | 74 (41.5) | 28 (37.8) | 46 (62.2) | 91 (51.1) | 28 (30.8) | 63 (69.2) | 0.60 |
| throw it away | 91 (51.1) | 30 (33.0) | 61 (67.0) | 73 (41.0) | 23 (31.5) | 50 (68.5) | 14 (7.9) | 8 (57.1) | 6 (42.9) | 0.16 |
| mix with another animals' milk | 2 (1.1) | 1 (50.0) | 1 (50.0) | 66 (37.1) | 24 (36.4) | 42 (63.6) | 110 (61.8) | 36 (32.7) | 74 (67.3) | 0.72~ |
| do not milk sheep and goat, but if cow or buffalo, treat and sell its milk/slaughter infected ruminant | 7 (3.9) | 2 (28.6) | 5 (71.4) | 1 (0.5) | 0 (0.0) | 1 (100.0) | 170 (90.8) | 59 (34.7) | 111 (65.3) | 0.90~ |

\*p-value is chi square test. The statistics was carried out in this table by row ~ ; P value for Fisher's exact test

**Table S8:** Practice of animal vaccination against brucellosis by cases and control

|  | Total**<br>266 own<br>animals |  | Cases (n=217)<br>106 own<br>animals |  | Control (434)<br>160 own animals |  | *p-value |
| --- | --- | --- | --- | --- | --- | --- | --- |
|  | No. | % | No. | % | No. | % |  |
| **Do you vaccinate animals against brucellosis (n=266)** |  |  |  |  |  |  |  |
| Yes | 90 | 33.8 | 11 | 10.4 | 79 | 49.4 | **0.0001** |
| No | 45 | 16.9 | 25 | 23.6 | 20 | 12.5 |  |
| I do not know | 109 | 41.0 | 62 | 58.5 | 47 | 29.4 |  |
| I vaccinate but I do not know what type of vaccine it is | 22 | 8.3 | 8 | 7.5 | 14 | 8.8 |  |
| **Who vaccinate the animals** |  |  |  |  |  |  |  |
| Gov vet free of charge | 111 | 94.9 | 21 | 91.3 | 90 | 95.7 | **0.40~** |
| Gov vet with charge | 5 | 4.3 | 2 | 8.7 | 3 | 3.2 |  |
| Private vet | 1 | 0.8 | 0 | 0.0 | 1 | 1.1 |  |
| **Do you pay for vaccinating animals against brucellosis** |  |  |  |  |  |  |  |
| Yes | 40 | 35.7 | 11 | 45.8 | 29 | 33.0 | **0.24** |
| No | 72 | 64.3 | 13 | 54.2 | 59 | 77.0 |  |
| **Will you vaccinate your animals in the future** |  |  |  |  |  |  |  |
| Yes (including those who actually practice animal vaccination) | 190 | 71.7 | 67 | 63.2 | 123 | 77.4 | **0.013** |
| No | 36 | 13.6 | 22 | 20.8 | 14 | 8.8 |  |
| I do not know | 39 | 14.7 | 17 | 16.0 | 22 | 13.8 |  |

\*p-value is for chi square test \*\* the total number for some questions does not equal 651 because of lack of response of some participants

**Table S9:**
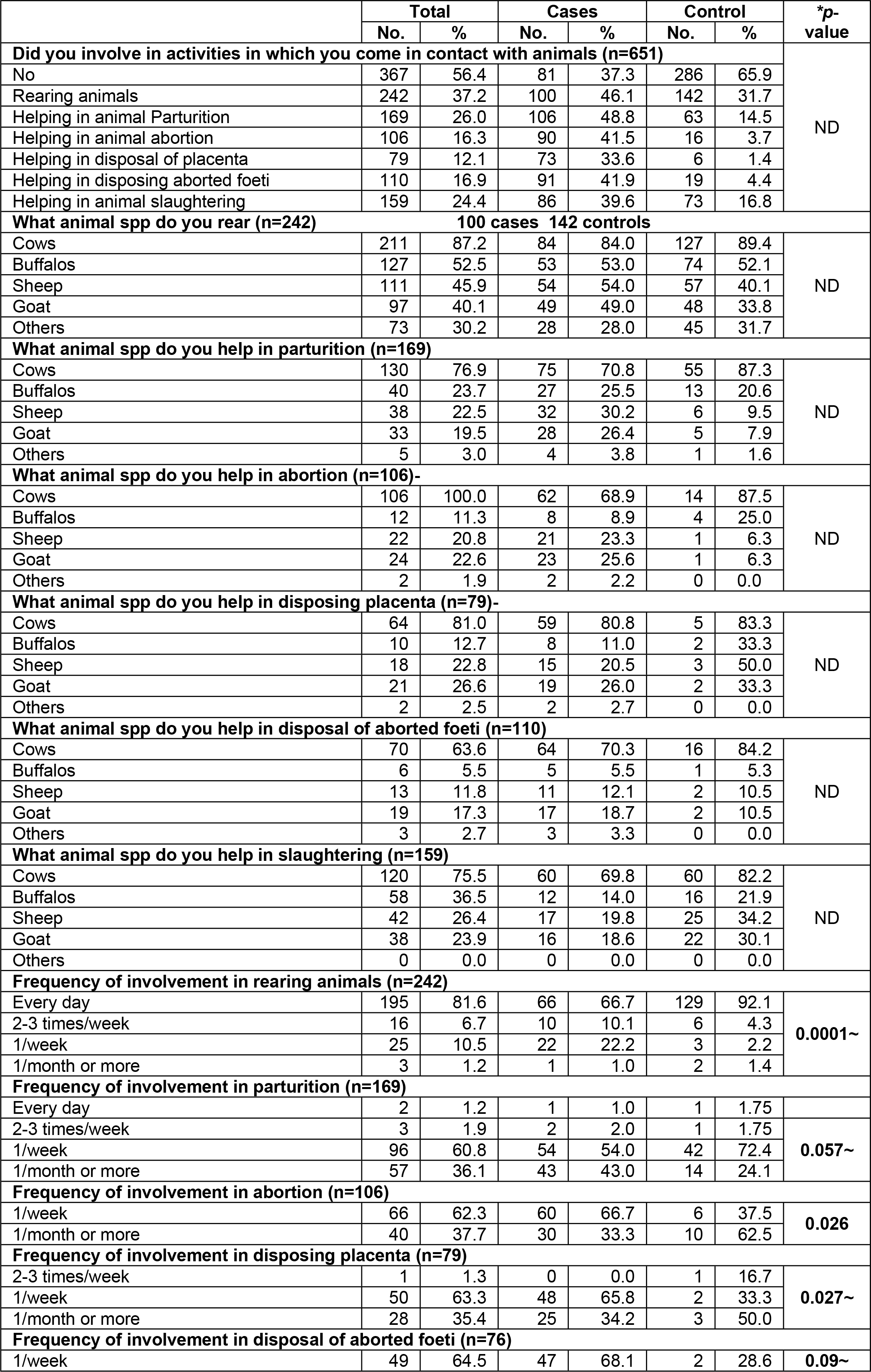
Practice of animal husbandry by cases and controls

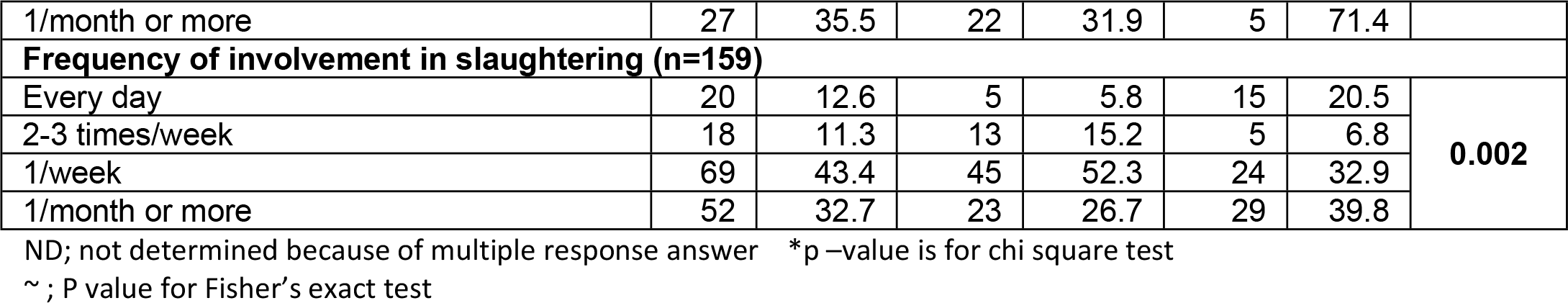

**Table S10:**
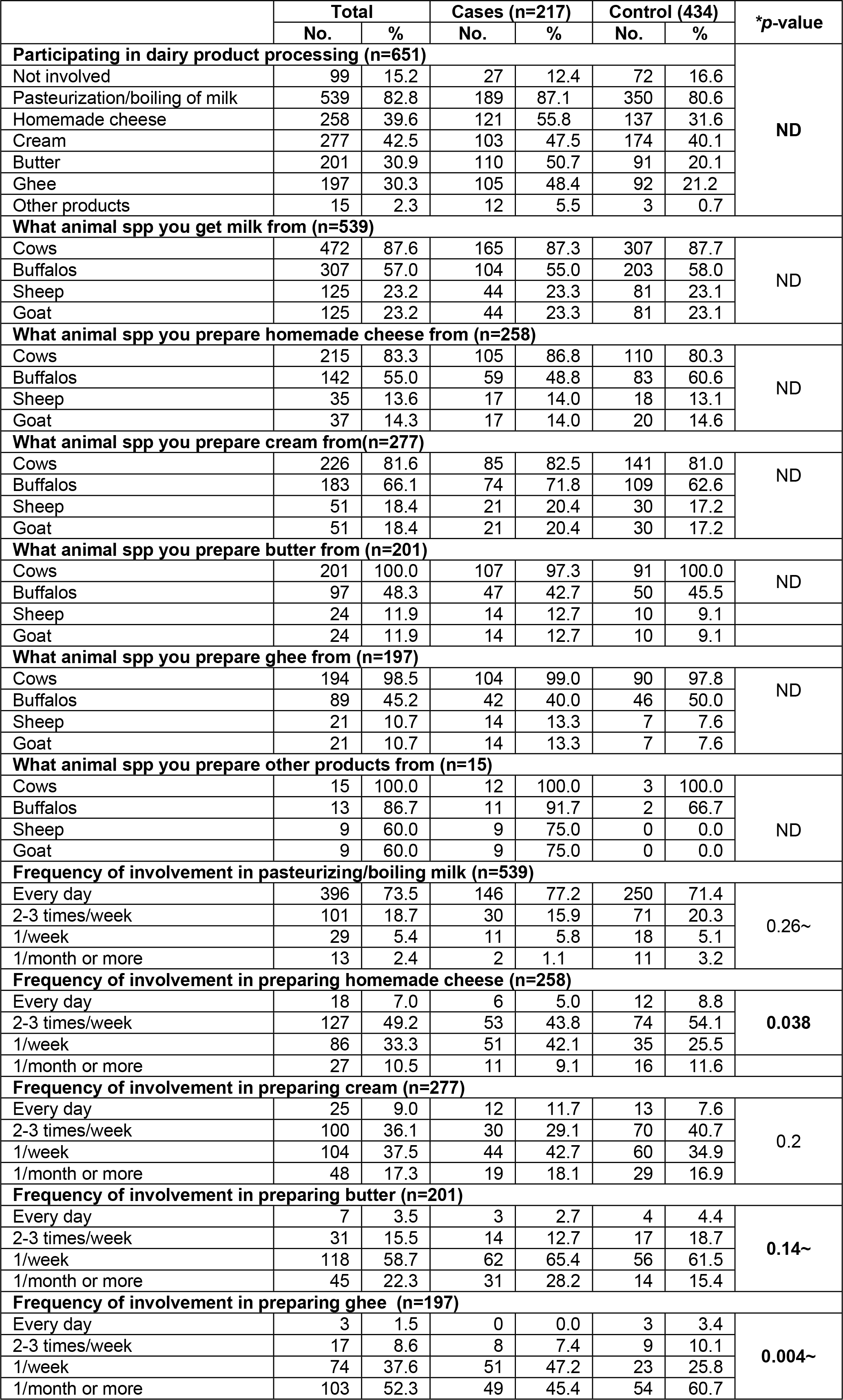
Practice of dairy farming by cases and controls

|  | Total |  | Cases (n=217) |  | Control (434) |  | *p-value |
| --- | --- | --- | --- | --- | --- | --- | --- |
|  | No. | % | No. | % | No. | % |  |
| Participating in dairy product processing (n=651) |  |  |  |  |  |  |  |
| Not involved | 99 | 15.2 | 27 | 12.4 | 72 | 16.6 | ND |
| Pasteurization/boiling of milk | 539 | 82.8 | 189 | 87.1 | 350 | 80.6 |  |
| Homemade cheese | 258 | 39.6 | 121 | 55.8 | 137 | 31.6 |  |
| Cream | 277 | 42.5 | 103 | 47.5 | 174 | 40.1 |  |
| Butter | 201 | 30.9 | 110 | 50.7 | 91 | 20.1 |  |
| Ghee | 197 | 30.3 | 105 | 48.4 | 92 | 21.2 |  |
| Other products | 15 | 2.3 | 12 | 5.5 | 3 | 0.7 |  |
| What animal spp you get milk from (n=539) |  |  |  |  |  |  |  |
| Cows | 472 | 87.6 | 165 | 87.3 | 307 | 87.7 | ND |
| Buffalos | 307 | 57.0 | 104 | 55.0 | 203 | 58.0 |  |
| Sheep | 125 | 23.2 | 44 | 23.3 | 81 | 23.1 |  |
| Goat | 125 | 23.2 | 44 | 23.3 | 81 | 23.1 |  |
| What animal spp you prepare homemade cheese from (n=258) |  |  |  |  |  |  |  |
| Cows | 215 | 83.3 | 105 | 86.8 | 110 | 80.3 | ND |
| Buffalos | 142 | 55.0 | 59 | 48.8 | 83 | 60.6 |  |
| Sheep | 35 | 13.6 | 17 | 14.0 | 18 | 13.1 |  |
| Goat | 37 | 14.3 | 17 | 14.0 | 20 | 14.6 |  |
| What animal spp you prepare cream from(n=277) |  |  |  |  |  |  |  |
| Cows | 226 | 81.6 | 85 | 82.5 | 141 | 81.0 | ND |
| Buffalos | 183 | 66.1 | 74 | 71.8 | 109 | 62.6 |  |
| Sheep | 51 | 18.4 | 21 | 20.4 | 30 | 17.2 |  |
| Goat | 51 | 18.4 | 21 | 20.4 | 30 | 17.2 |  |
| What animal spp you prepare butter from (n=201) |  |  |  |  |  |  |  |
| Cows | 201 | 100.0 | 107 | 97.3 | 91 | 100.0 | ND |
| Buffalos | 97 | 48.3 | 47 | 42.7 | 50 | 45.5 |  |
| Sheep | 24 | 11.9 | 14 | 12.7 | 10 | 9.1 |  |
| Goat | 24 | 11.9 | 14 | 12.7 | 10 | 9.1 |  |
| What animal spp you prepare ghee from (n=197) |  |  |  |  |  |  |  |
| Cows | 194 | 98.5 | 104 | 99.0 | 90 | 97.8 | ND |
| Buffalos | 89 | 45.2 | 42 | 40.0 | 46 | 50.0 |  |
| Sheep | 21 | 10.7 | 14 | 13.3 | 7 | 7.6 |  |
| Goat | 21 | 10.7 | 14 | 13.3 | 7 | 7.6 |  |
| What animal spp you prepare other products from (n=15) |  |  |  |  |  |  |  |
| Cows | 15 | 100.0 | 12 | 100.0 | 3 | 100.0 | ND |
| Buffalos | 13 | 86.7 | 11 | 91.7 | 2 | 66.7 |  |
| Sheep | 9 | 60.0 | 9 | 75.0 | 0 | 0.0 |  |
| Goat | 9 | 60.0 | 9 | 75.0 | 0 | 0.0 |  |
| Frequency of involvement in pasteurizing/boiling milk (n=539) |  |  |  |  |  |  |  |
| Every day | 396 | 73.5 | 146 | 77.2 | 250 | 71.4 | 0.26~ |
| 2-3 times/week | 101 | 18.7 | 30 | 15.9 | 71 | 20.3 |  |
| 1/week | 29 | 5.4 | 11 | 5.8 | 18 | 5.1 |  |
| 1/month or more | 13 | 2.4 | 2 | 1.1 | 11 | 3.2 |  |
| Frequency of involvement in preparing homemade cheese (n=258) |  |  |  |  |  |  |  |
| Every day | 18 | 7.0 | 6 | 5.0 | 12 | 8.8 | 0.038 |
| 2-3 times/week | 127 | 49.2 | 53 | 43.8 | 74 | 54.1 |  |
| 1/week | 86 | 33.3 | 51 | 42.1 | 35 | 25.5 |  |
| 1/month or more | 27 | 10.5 | 11 | 9.1 | 16 | 11.6 |  |
| Frequency of involvement in preparing cream (n=277) |  |  |  |  |  |  |  |
| Every day | 25 | 9.0 | 12 | 11.7 | 13 | 7.6 | 0.2 |
| 2-3 times/week | 100 | 36.1 | 30 | 29.1 | 70 | 40.7 |  |
| 1/week | 104 | 37.5 | 44 | 42.7 | 60 | 34.9 |  |
| 1/month or more | 48 | 17.3 | 19 | 18.1 | 29 | 16.9 |  |
| Frequency of involvement in preparing butter (n=201) |  |  |  |  |  |  |  |
| Every day | 7 | 3.5 | 3 | 2.7 | 4 | 4.4 | 0.14~ |
| 2-3 times/week | 31 | 15.5 | 14 | 12.7 | 17 | 18.7 |  |
| 1/week | 118 | 58.7 | 62 | 65.4 | 56 | 61.5 |  |
| 1/month or more | 45 | 22.3 | 31 | 28.2 | 14 | 15.4 |  |
| Frequency of involvement in preparing ghee (n=197) |  |  |  |  |  |  |  |
| Every day | 3 | 1.5 | 0 | 0.0 | 3 | 3.4 | 0.004~ |
| 2-3 times/week | 17 | 8.6 | 8 | 7.4 | 9 | 10.1 |  |
| 1/week | 74 | 37.6 | 51 | 47.2 | 23 | 25.8 |  |
| 1/month or more | 103 | 52.3 | 49 | 45.4 | 54 | 60.7 |  |

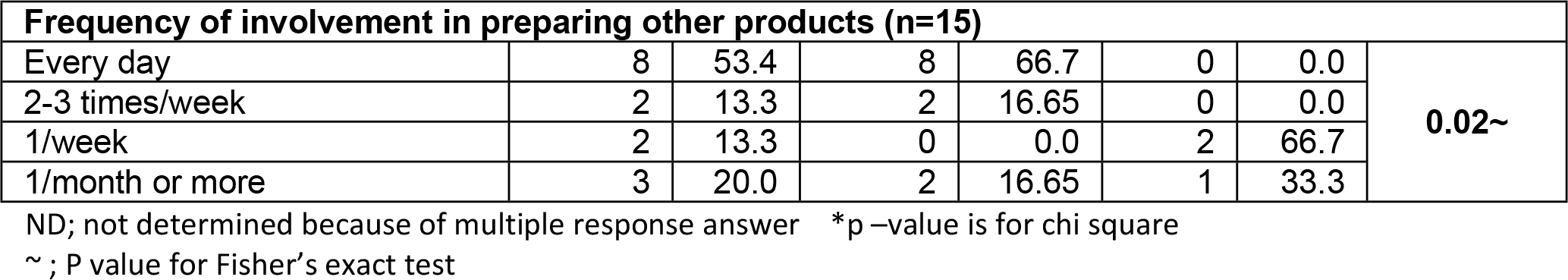

**Table S11:** involvement of cases and controls in buying/selling livestock products (milk, dairy products)

|  | Usually |  |  | Sometimes |  |  | Never |  |  | *p-value |
| --- | --- | --- | --- | --- | --- | --- | --- | --- | --- | --- |
|  | Total | Cases | Control | Total | Cases | Control | Total | Cases | Control |  |
|  | No. (%) | No. (%) | No. (%) | No. (%) | No. (%) | No. (%) | No. (%) | No. (%) | No. (%) |  |
| Buy/sell livestock products (milk, dairy products) |  |  |  |  |  |  |  |  |  |  |
| in the market | 432 (66.4) | 139 (32.2) | 293 (67.8) | 77 (11.4) | 18 (24.3) | 59 (75.7) | 145 (22.3) | 60 (41.4) | 85 (58.6) | 0.018 |
| in the same village | 538 (82.6) | 190 (35.3) | 348 (64.7) | 84 (12.9) | 21 (25.0) | 63 (75.0) | 29 (4.5) | 6 (20.7) | 23 (79.3) | 0.059 |
| in the neighboring village | 40 (6.1) | 19 (47.5) | 21 (52.5) | 332 (51.0) | 91 (27.4) | 241 (72.6) | 279 (42.9) | 107 (38.4) | 172 (61.6) | 0.002 |
| in far villages within the same district | 15 (2.3) | 6 (40.0) | 9 (60.0) | 132 (20.3) | 42 (31.8) | 90 (68.2) | 504 (77.4) | 169 (33.5) | 335 (66.5) | 0.788 |
| outside district but within governorate | 14 (2.2) | 4 (28.6) | 10 (71.4) | 107 (16.4) | 28 (26.2) | 79 (73.8) | 530 (81.4) | 185 (34.9) | 345 (65.1) | 0.200~ |
| in other governorates | 5 (0.8) | 2 (40.0) | 3 (60.0) | 69 (10.6) | 16 (23.2) | 53 (76.8) | 577 (88.6) | 199 (34.5) | 378 (65.5) | 0.144~ |

\*p-value is for chi square test The statistics was carried out in this table by row ~ ; P value for Fisher's exact test

**Table S12:**
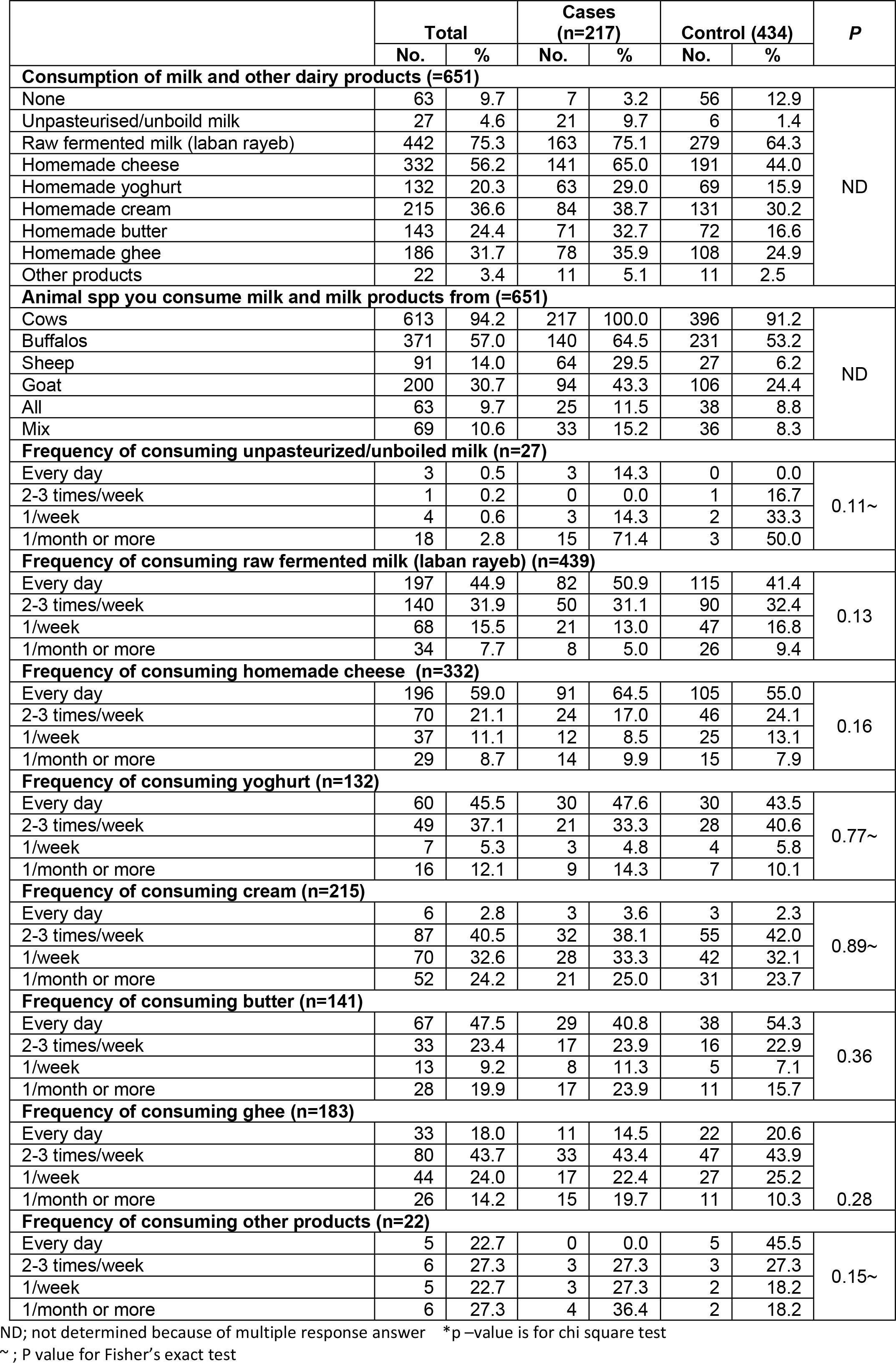
Consumption of milk and milk products by cases and controls

**Table S13:**
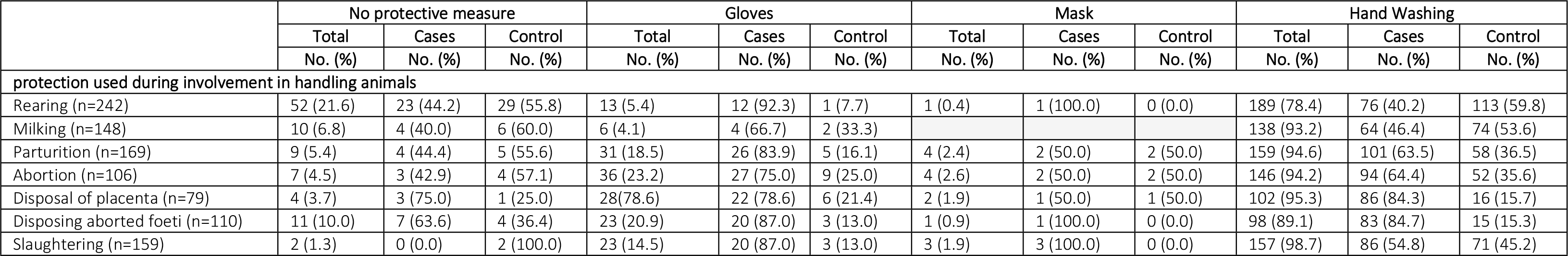
Protective measures applied by cases and controls during the practice of animal husbandry

|  | No protective measure |  |  | Gloves |  |  | Mask |  |  | Hand Washing |  |  |
| --- | --- | --- | --- | --- | --- | --- | --- | --- | --- | --- | --- | --- |
|  | Total | Cases | Control | Total | Cases | Control | Total | Cases | Control | Total | Cases | Control |
|  | No. (%) | No. (%) | No. (%) | No. (%) | No. (%) | No. (%) | No. (%) | No. (%) | No. (%) | No. (%) | No. (%) | No. (%) |
| **protection used during involvement in handling animals** |  |  |  |  |  |  |  |  |  |  |  |  |
| Rearing (n=242) | 52 (21.6) | 23 (44.2) | 29 (55.8) | 13 (5.4) | 12 (92.3) | 1 (7.7) | 1 (0.4) | 1 (100.0) | 0 (0.0) | 189 (78.4) | 76 (40.2) | 113 (59.8) |
| Milking (n=148) | 10 (6.8) | 4 (40.0) | 6 (60.0) | 6 (4.1) | 4 (66.7) | 2 (33.3) |  |  |  | 138 (93.2) | 64 (46.4) | 74 (53.6) |
| Parturition (n=169) | 9 (5.4) | 4 (44.4) | 5 (55.6) | 31 (18.5) | 26 (83.9) | 5 (16.1) | 4 (2.4) | 2 (50.0) | 2 (50.0) | 159 (94.6) | 101 (63.5) | 58 (36.5) |
| Abortion (n=106) | 7 (4.5) | 3 (42.9) | 4 (57.1) | 36 (23.2) | 27 (75.0) | 9 (25.0) | 4 (2.6) | 2 (50.0) | 2 (50.0) | 146 (94.2) | 94 (64.4) | 52 (35.6) |
| Disposal of placenta (n=79) | 4 (3.7) | 3 (75.0) | 1 (25.0) | 28 (78.6) | 22 (78.6) | 6 (21.4) | 2 (1.9) | 1 (50.0) | 1 (50.0) | 102 (95.3) | 86 (84.3) | 16 (15.7) |
| Disposing aborted foeti (n=110) | 11 (10.0) | 7 (63.6) | 4 (36.4) | 23 (20.9) | 20 (87.0) | 3 (13.0) | 1 (0.9) | 1 (100.0) | 0 (0.0) | 98 (89.1) | 83 (84.7) | 15 (15.3) |
| Slaughtering (n=159) | 2 (1.3) | 0 (0.0) | 2 (100.0) | 23 (14.5) | 20 (87.0) | 3 (13.0) | 3 (1.9) | 3 (100.0) | 0 (0.0) | 157 (98.7) | 86 (54.8) | 71 (45.2) |

**Table S14:**
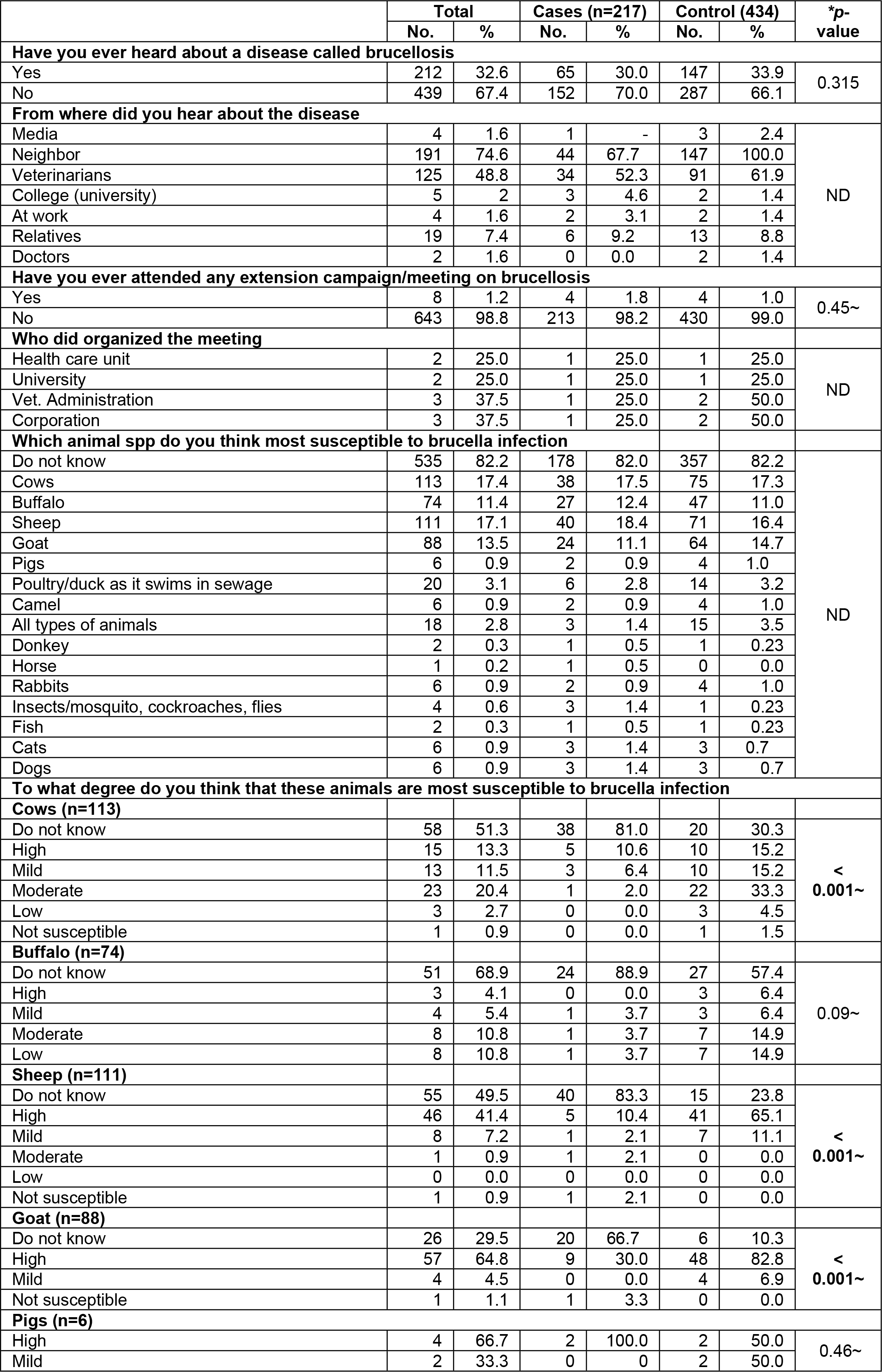
knowledge and perceived animal susceptibility regarding brucellosis as reported by cases and controls

|  |  |  |  |  |  |  |  |
| --- | --- | --- | --- | --- | --- | --- | --- |
| Poultry/duck as it swims in sewage (n=20) |  |  |  |  |  |  |  |
| Do not know | 3 | 15.0 | 3 | 50.0 | 0 | 0.0 | 0.018~ |
| High | 16 | 80.0 | 3 | 50.0 | 13 | 92.9 |  |
| Mild | 1 | 5.0 | 0 | 0.0 | 1 | 7.1 |  |
| Camel (n=6) |  |  |  |  |  |  |  |
| High | 2 | 33.3 | 1 | 50.0 | 1 | 25.0 | 0.73~ |
| Mild | 2 | 33.3 | 0 | 0.0 | 2 | 50.0 |  |
| Moderate | 1 | 16.7 | 0 | 0.0 | 1 | 50.0 |  |
| Low | 1 | 16.7 | 1 | 50.0 | 0 | 0.0 |  |
| All types of animals (n=18) |  |  |  |  |  |  |  |
| High | 18 | 100.0 | 3 | 100.0 | 13 | 100.0 | - |
| Donkey (n=2) |  |  |  |  |  |  |  |
| High | 1 | 50.0 | 1 | 100.0 | 0 | 0.0 | 0.99~ |
| Low | 1 | 50.0 | 0 | 0.0 | 1 | 100.0 |  |
| Horse (n=1) |  |  |  |  |  |  |  |
| High | 1 | 100.0 | 1 | 100.0 | 0 | 100.0 | ND |
| Rabbits (n=6) |  |  |  |  |  |  |  |
| Do not know | 1 | 16.6 | 1 | 50.0 | 0 | 0.0 | 0.99~ |
| High | 3 | 50.0 | 1 | 50.0 | 2 | 50.0 |  |
| Mild | 1 | 16.7 | 0 | 0.0 | 1 | 25.0 |  |
| Moderate | 1 | 16.7 | 0 | 0.0 | 1 | 25.0 |  |
| Insects/mosquito, cockroaches, flies (n=4) |  |  |  |  |  |  |  |
| Do not know | 2 | 50.0 | 2 | 66.7 | 0 | 0.0 | 0.99~ |
| High | 2 | 50.0 | 1 | 33.3 | 1 | 100.0 |  |
| Fish (n=2) |  |  |  |  |  |  |  |
| High | 1 | 50.0 | 0 | 0.0 | 1 | 100.0 | 0.99~ |
| Mild | 1 | 50.0 | 1 | 100.0 | 0 | 0.0 |  |
| Cats (n=6) |  |  |  |  |  |  |  |
| High | 6 | 100.0 | 3 | 100.0 | 3 | 100.0 | ND |
| Dogs (n=6) |  |  |  |  |  |  |  |
| High | 6 | 100.0 | 3 | 100.0 | 3 | 100.0 | ND |
| What age do you think that these animals are most susceptible to brucella infection |  |  |  |  |  |  |  |
| Cows (n=113) |  |  |  |  |  |  |  |
| Do not know | 67 | 59.3 | 30 | 48.4 | 37 | 37.8 | 0.24~ |
| All ages | 20 | 17.7 | 24 | 38.7 | 42 | 42.6 |  |
| < 1 year | 23 | 20.4 | 6 | 9.7 | 18 | 18.4 |  |
| > 1 year | 3 | 2.6 | 2 | 3.2 | 1 | 1.2 |  |
| Buffalo (n=74) |  |  |  |  |  |  |  |
| Do not know | 57 | 77.0 | 30 | 56.6 | 27 | 40.3 | 0.03~ |
| All ages | 8 | 10.8 | 22 | 41.5 | 32 | 47.8 |  |
| < 1 year | 7 | 9.5 | 0 | 0 | 7 | 10.4 |  |
| Large ages | 2 | 2.7 | 1 | 1.9 | 1 | 1.5 |  |
| Sheep (n=111) |  |  |  |  |  |  |  |
| Do not know | 20 | 18.0 | 4 | 10.0 | 16 | 22.2 | 0.37~ |
| All ages | 56 | 50.5 | 21 | 32.5 | 35 | 48.6 |  |
| < 1 year | 34 | 30.6 | 14 | 35.0 | 20 | 27.8 |  |
| Large ages | 1 | 0.9 | 1 | 2.5 | 1 | 1.4 |  |
| Goat (n=88) |  |  |  |  |  |  |  |
| Do not know | 46 | 52.3 | 20 | 48.8 | 26 | 35.1 | 0.008~ |
| All ages | 24 | 27.3 | 19 | 46.4 | 32 | 43.2 |  |
| < 1 year | 17 | 19.3 | 1 | 2.4 | 16 | 21.7 |  |
| Large ages | 1 | 1.1 | 1 | 2.4 | 0 | 0 |  |
| Pigs (n=6) |  |  |  |  |  |  |  |
| Do not know | 6 | 100.0 | 2 | 100.0 | 4 | 100.0 | ND |
| Poultry/duck as it swims in sewage (n=20) |  |  |  |  |  |  |  |
| Do not know | 17 | 85.0 | 6 | 100.0 | 11 | 78.6 | 0.52~ |
| All ages | 3 | 15.0 | 0 | 0.0 | 3 | 21.4 |  |
| Camel (n=6) |  |  |  |  |  |  |  |
| All ages | 4 | 66.7 | 2 | 100.0 | 2 | 50.0 | 0.46~ |
| < 1 year | 2 | 33.3 | 0 | 0.0 | 2 | 50.0 |  |
| All types of animals (n=18) |  |  |  |  |  |  |  |
| Do not know | 6 | 33.3 | 2 | 40.0 | 4 | 30.8 | 0.99~ |
| All ages | 12 | 66.7 | 3 | 60.0 | 9 | 69.2 |  |
| Donkey (n=2) |  |  |  |  |  |  |  |
| Do not know | 1 | 50.0 | 1 | 100.0 | 0 | 0.0 | 0.99~ |
| < 1 year | 1 | 50.0 | 0 | 0.0 | 1 | 100.0 |  |
| Horse (n=1) |  |  |  |  |  |  |  |
| Do not know | 1 | 100.0 | 1 | 100.0 | 0 | 100.0 | ND |
| Rabbits (n=6) |  |  |  |  |  |  |  |
| Do not know | 6 | 100.0 | 2 | 100.0 | 4 | 100.0 | ND |
| Insects/mosquito, cockroaches, flies (n=4) |  |  |  |  |  |  |  |
| Do not know | 4 | 100.0 | 3 | 100.0 | 1 | 100.0 | ND |
| **Fish (n=2)** |  |  |  |  |  |  |  |
| Do not know | 1 | 50.0 | 1 | 100.0 | 0 | 0.0 | 0.99~ |
| All ages | 1 | 50.0 | 0 | 0.0 | 1 | 100.0 |  |
| **Cats (n=6)** |  |  |  |  |  |  |  |
| Do not know | 1 | 16.7 | 0 | 0.0 | 1 | 33.3 | 0.99~ |
| All ages | 5 | 83.3 | 3 | 100.0 | 2 | 66.7 |  |
| **Dogs (n=6)** |  |  |  |  |  |  |  |
| Do not know | 1 | 16.7 | 0 | 0.0 | 1 | 33.3 | 0.99~ |
| All ages | 5 | 83.3 | 3 | 100.3 | 2 | 66.7 |  |

ND; not determined because of multiple response answer \*p –value is for chi square test
~ ; P value for Fisher's exact test

**Table S15:**
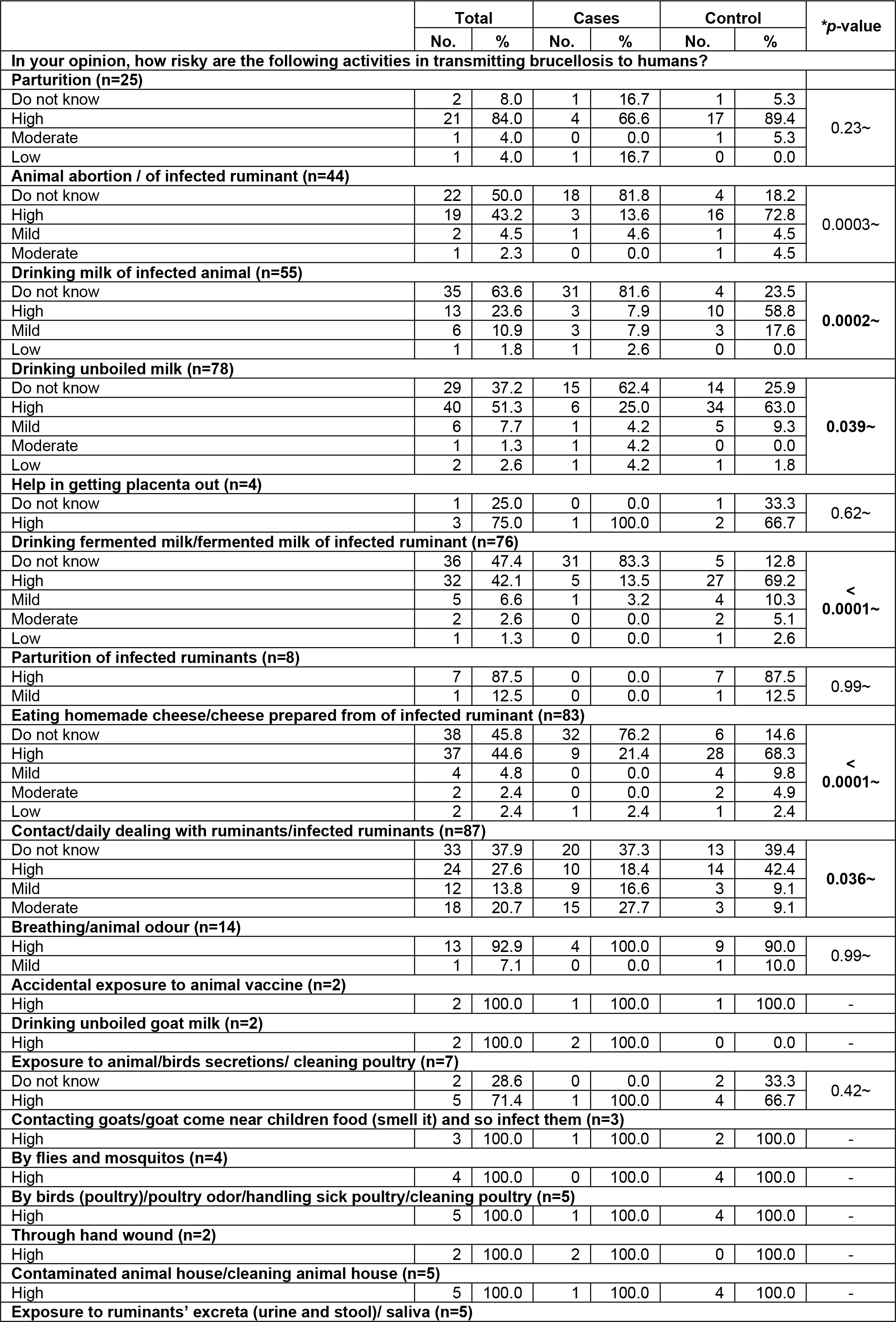
Perception of risky exposures by cases and controls

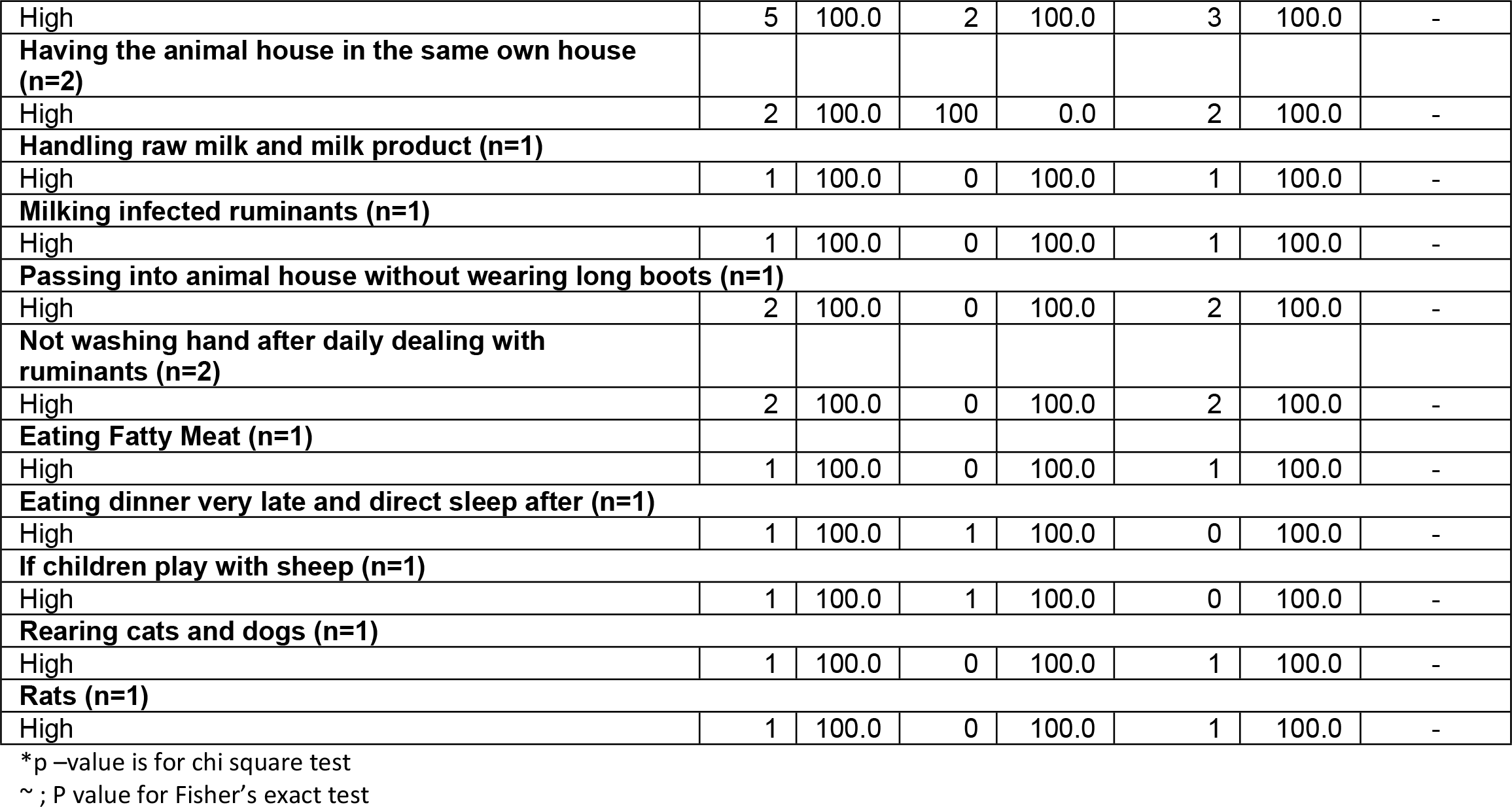

